# Germline loss of *MBD4* predisposes to leukaemia due to a mutagenic cascade driven by 5mC

**DOI:** 10.1101/180588

**Authors:** Mathijs A. Sanders, Edward Chew, Christoffer Flensburg, Annelieke Zeilemaker, Sarah E. Miller, Adil S. al Hinai, Ashish Bajel, Bram Luiken, Melissa Rijken, Tamara Mclennan, Remco M. Hoogenboezem, François G. Kavelaars, Marnie E. Blewitt, Eric M. Bindels, Warren S. Alexander, Bob Löwenberg, Andrew W. Roberts, Peter J.M. Valk, Ian J. Majewski

## Abstract

Cytosine methylation is essential for normal mammalian development, yet also provides a major mutagenic stimulus. Methylcytosine (5mC) is prone to spontaneous deamination, which introduces cytosine to thymine transition mutations (C>T) upon replication^1^. Cells endure hundreds of 5mC deamination events each day and an intricate repair network is engaged to restrict this damage. Central to this network are the DNA glycosylases MBD4^2^ and TDG^3^,^4^, which recognise T:G mispairing and initiate base excision repair (BER). Here we describe a novel cancer predisposition syndrome resulting from germline biallelic inactivation of MBD4 that leads to the development of acute myeloid leukaemia (AML). These leukaemias have an extremely high burden of C>T mutations, specifically in the context of methylated CG dinucleotides (CG>TG). This dependence on 5mC as a source of mutations may explain the remarkable observation that MBD4-deficient AMLs share a common set of driver mutations, including biallelic mutations in *DNMT3A* and hotspot mutations in *IDH1/IDH2*. By assessing serial samples taken over the course of treatment, we highlight a critical interaction with somatic mutations in *DNMT3A* that accelerates leukaemogenesis and accounts for the conserved path to AML. MBD4-deficiency was also detected, rarely, in sporadic cancers, which display the same mutational signature. Collectively these cancers provide a model of 5mC-dependent hypermutation and reveal factors that shape its mutagenic influence.

---

We identified three patients with AML, including two siblings, that were distinctive because of their early age of onset (all <35 years old) and an extremely high mutational burden (~33-fold above what is typical for AML) **(Fig. 1a, Clinical Synopsis)**. Virtually all of the somatic mutations identified were C>T in the context of a CG dinucleotide (>95% of SNVs) **(Fig. 1b, Extended Data Fig. 1)**. This differs markedly from the distribution of C>T mutations in AML generally and is more refined than the mutational signature ascribed to ageing, which includes a strong contribution from 5mC deamination^5^. All three cases carried rare germline loss-of-function variants in the gene encoding the DNA glycosylase MBD4^2^ **(Fig. 1c, Extended Data Table 1)**. Case EMC-AML-1 carried a homozygous *MBD4* in-frame deletion of Histidine 567 (His567) in the glycosylase domain. An *in vitro* glycosylase assay confirmed that loss of His567 results in a catalytically inactive MBD4 protein **(Fig. 1c)**. The siblings (WEHI-AML-1, WEHI-AML-2) were compound heterozygotes with a frameshift in exon 3 and a variant that disrupts the splice acceptor of exon 7 **(Fig. 1c, Extended Data Table 1)**. Analysis of the *MBD4* mRNA allowed for phasing of the variants to distinct alleles and confirmed aberrant splicing that excludes exon 7 and disrupts the glycosylase domain **(Extended Data Fig. 2)**. MBD4 has not previously been associated with haematological malignancy, but somatic mutations have been detected in sporadic colon cancers with mismatch repair (MMR) deficiency^6^,^7^. Two patients (EMC-AML-1, WEHI-AML-2) also had colorectal polyps, a common manifestation of DNA repair defects, including those associated with loss of BER components MUTYH^8^–^10^ and NTHL1^11^.

**Figure 1:**
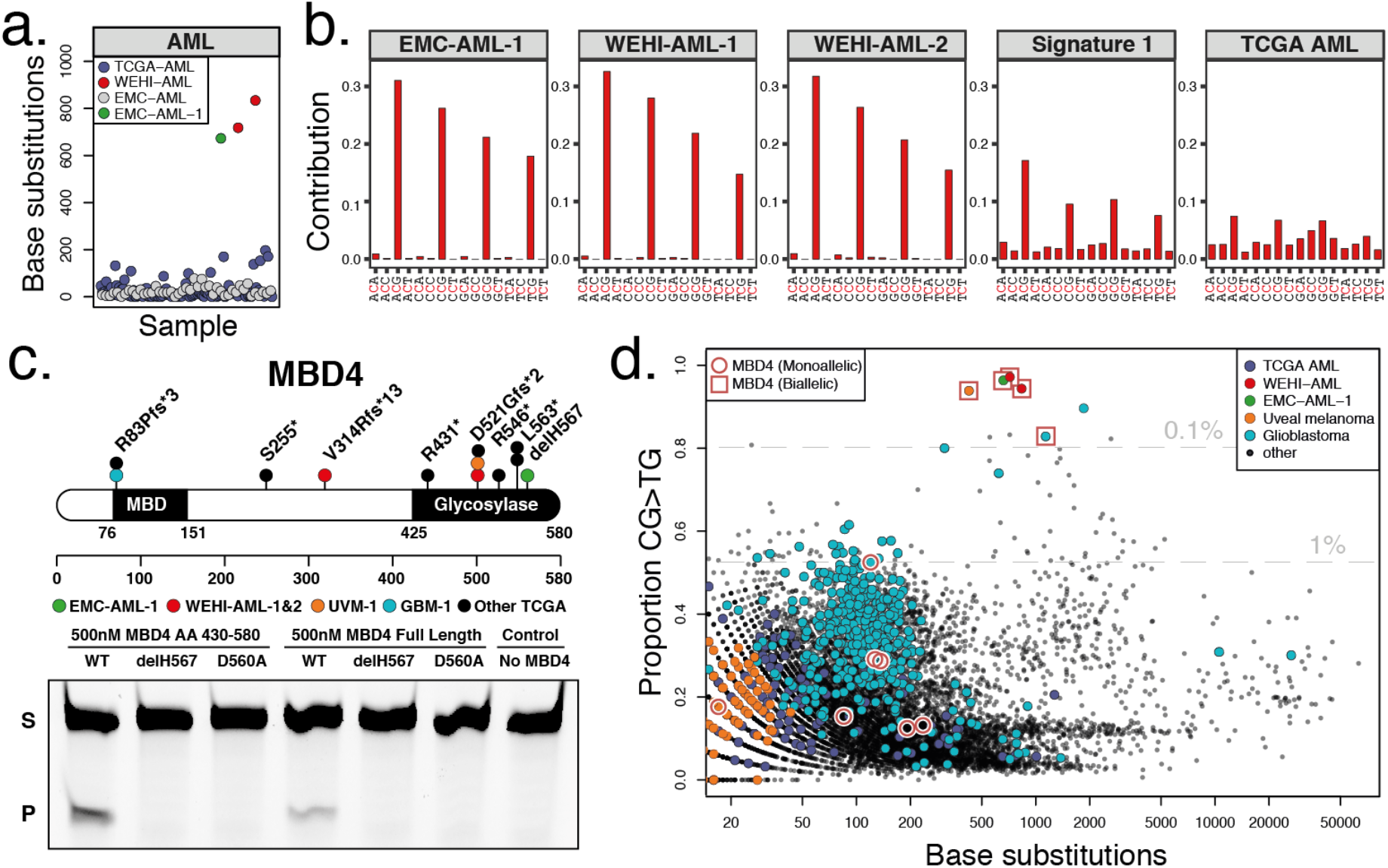
MBD4-deficient cancers exhibit a distinctive mutational signature. **a**, Mutation burden in AML, presented as number of base substitutions per exome. Data sourced from dbGAP, cases presented in order based on patient identifier (EMC: phs001027^27^ and TCGA: phs000178^28^). **b**, Trimer context of C>T mutations in three MBD4-deficient AML cases. For comparison we show Signature 1, the established signature associated with 5mC deamination, and all C>T mutations present in TCGA-AML. **c**, Schematic representation of MBD4, highlighting germline loss-of-function variants detected in the AML cases and in TCGA (at top). A glycosylase assay was performed to assess the activity of recombinant MBD4 (either AA430-580 or full length) - wildtype (WT), delH567, or the catalytically inactive mutant D560A^29^. Substrate (S) and Product (P). Consistent results were obtained in 5 experiments for MBD4 AA430-580 and 3 experiments for full length. **d**, Mutation profiles are shown for TCGA samples, focused on the proportion of CG>TG mutations. Samples with germline loss-of-function variants were designated either heterozygous (monoallelic) or completely inactivated (biallelic) based on the genotype of the cancer (includes somatic mutations). Grey lines mark the top 1% and 0.1% of all cases and select tumour types are highlighted.

**Figure 2:**
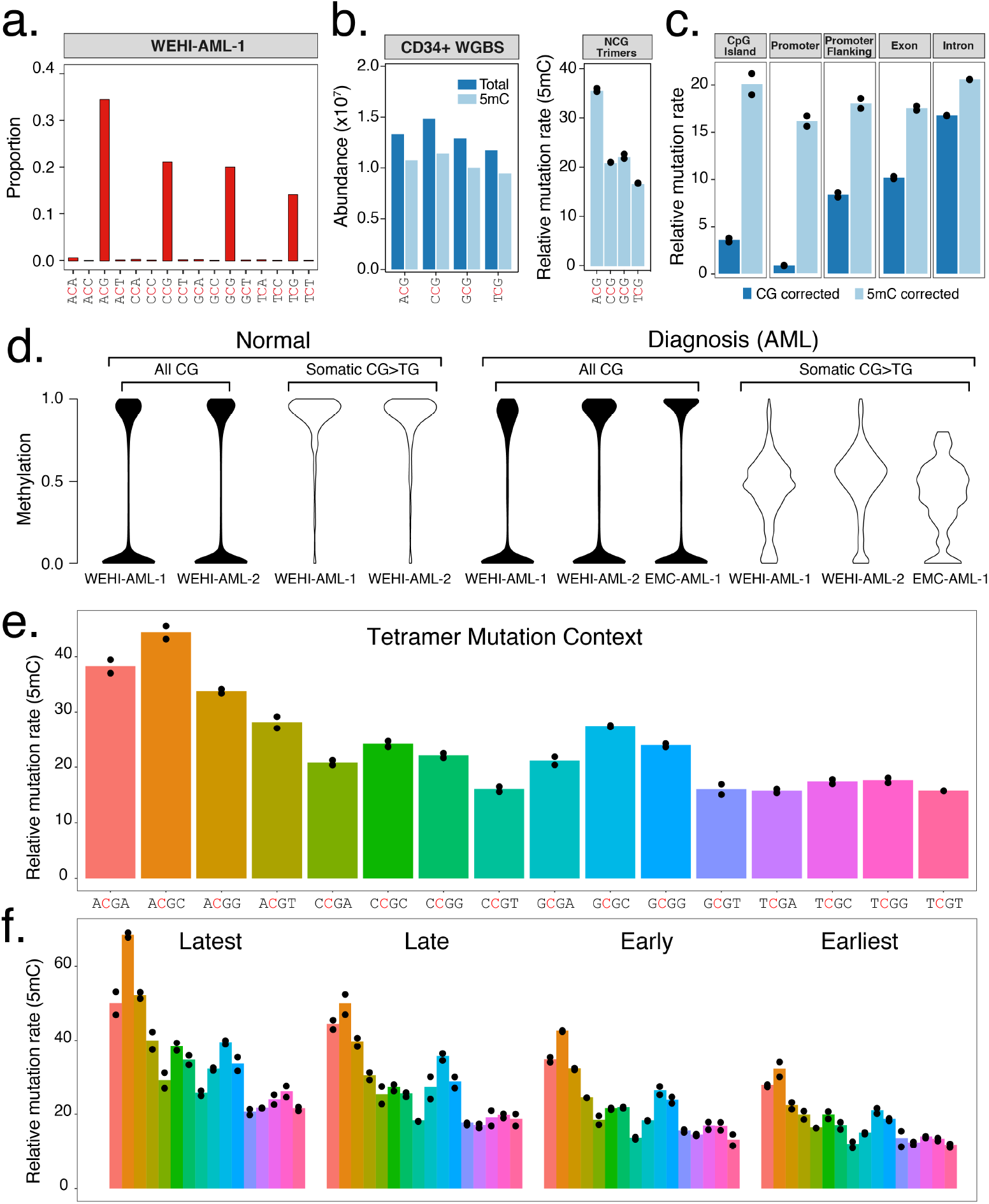
Damage due to 5mC deamination is influenced by genetic and epigenetic features. **a**, Trimer context of C>T mutations identified in whole genome sequencing from WEHI-AML-1 (14,313 mutations). Full signatures for both genomes are provided **(Extended Data Fig. 1b)**. **b**, Abundance and methylation status for NCG trimers from WGBS data from CD34+ cells^30^. A relative mutation rate was calculated for each trimer, accounting for differences in abundance and 5mC status in CD34+ cells and scaled to account for total mutation load (see Methods). Individual values are plotted (n=2) and the bar shows the mean. **c**, Relative mutation rate in different genomic features per Mb of CG dinucleotides (CG corrected), or corrected for methylation status in CD34+ cells (5mC corrected). **d**, 5mC assessed by RRBS. Cytosine levels were assessed at CG dinucleotides following bisulfite conversion, either globally (all CG) or at sites of somatic mutations. Normal bone marrow was at remission. **e**, The relative mutation rate was calculated for NCGN tetramers in WEHI-AML-1 and WEHI-AML-2, then separated by replication timing (**f**) (n=2, as above).

We accessed large cancer databases to explore the link between MBD4-deficiency and the distinctive CG>TG signature. Analysis of the Cancer Genome Atlas (TCGA) identified nine cases, from 10,683 total, that carried germline loss-of-function variants in *MBD4* **(Fig. 1c, Extended Data Table 1)**. In two cases, a uveal melanoma (TCGA-UVM-1) and a glioblastoma multiforme (TCGA-GBM-1), splice site mutations were accompanied by loss of the wildtype *MBD4* allele due to somatic copy number alterations **(Extended Data Fig. 3a)**. Analysis of RNA sequencing from both tumours confirmed aberrant splicing of *MBD4*, predicted to result in protein truncation and loss of function **(Extended Data Fig. 3b)**. Both cases exhibited an elevated mutation rate and strong enrichment for CG>TG mutations **(Fig. 1d, Extended Data Fig. 1a)**. This signature was also observed in a glioma cell line, SW1783, that carries a homozygous truncating variant in *MBD4* at Leu563 **(Extended Data Fig. 1a).** The cancers that retained a wildtype allele did not display a prominent CG>TG signature **(Fig. 1d)**. These results suggest both alleles of *MBD4* must be inactivated to block its repair activity, which is consistent with other BER-associated cancer syndromes^8^,^11^. Analysis of a larger cohort will be required to determine whether heterozygous loss of *MBD4* predisposes to cancer.

**Figure 3:**
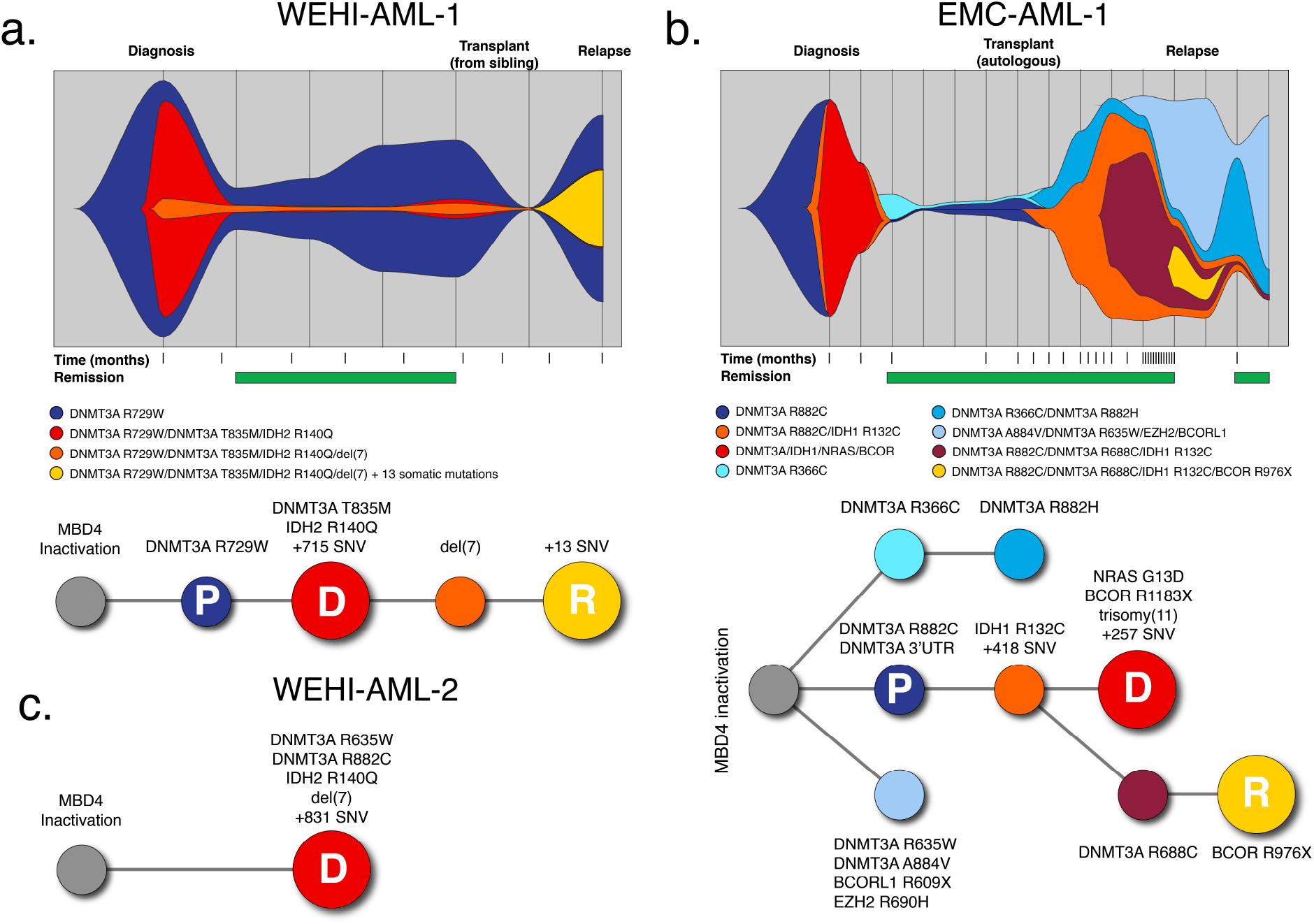
Germline MBD4-deficient patients share a common path to AML. Clonal evolution observed in WEHI-AML-1 (**a**) and EMC-AML-1 (**b**), based on variant allele frequencies derived from exomes and RNA-seq. For EMC-AML-1 single cell genotyping was used to resolve the clonal relationships. Each coloured area is proportional to the representation of the clone and vertical lines indicate sampling points^31^. Both patients experienced clonal haematopoiesis during remission. The transplant for WEHI-AML-1 was provided by WEHI-AML-2, which occurred four years prior to her own diagnosis of AML. Phylogenetic tree diagrams are shown to highlight the acquisition of key mutations for WEHI-AML-1 (**a**), EMC-AML-1 (**b**) and WEHI-AML-2 (**c**). The pre-malignant clone (P, in dark blue), and the AML clones evident at diagnosis (D, in red) and relapse (R, in yellow), are designated. Additional mutations were detected in exome data from EMC-AML-1 at relapse, but because of the presence of multiple clones they could not be unambiguously assigned.

Whole genome sequencing and methylation profiling were performed to refine the mutational signature associated with MBD4-deficiency in AML. While MBD4 is known to interact with the MMR pathway^12^, MBD4-deficienct leukaemias were largely devoid of small insertions and deletions, suggesting MMR remains intact. Overall, >15,000 substitution mutations were identified in each AML genome, of which >90% were CG>TG **(Fig. 2a, Extended Data Fig. 1b)**. The proportion of mutations was higher in the context of the ACG triplet and lower in the context of TCG, with CCG and GCG being intermediate. This difference remained after correction for trimer abundance and methylation status **(Fig. 2b)**, and was found to be significant in the exome data from the five MBD4-deficient cancers (p= 0.007937, Mann-Whitney U test) **(Extended Data Fig. 1)**. The ACA trimer was the most commonly mutated site outside of a CG context, and this matches the most common site of non-CG methylation^13^. The mutation rate for a given region was linked to 5mC abundance. Sparsely methylated regions, such as promoters and CG islands, were rarely mutated **(Fig. 2c)**. Correcting for 5mC abundance revealed a consistent mutation rate across different genomic features **(Fig. 2c)**. Reduced representation bisulfite sequencing (RRBS) confirmed that >95% of CG sites mutated in the AML were fully methylated in matched normal bone marrow available for two cases **(Fig. 2d)**. Assessment of the mutated sites in each AML directly revealed ~50% methylation, indicating the non-mutated CG site on the alternate allele was methylated **(Fig. 2d)**. Similar results were obtained when we assessed sites mutated in the MBD4-deficient glioblastoma **(Extended Data Fig. 4)**. We next assessed the influence of genetic and epigenetic features known to influence mutation rate^14^. Extending the analysis of sequence context to include one base either side of the CG identified higher mutation rates in the context of a 3’ cytosine (NCGC), with the highest rate at ACGC **(Fig. 2e)**. The relative mutation rate was not influenced by the transcriptional strand **(Extended Data Fig. 5a),** but was higher in late replicating regions **(Fig. 2f)** and at lowly expressed genes **(Extended Data Fig. 5b)**. Collectively these results suggest that while 5mC is the dominant factor contributing to the mutation rate, the local sequence context, replication timing and expression status also contribute. The differences between tetramers and enrichment in late replicating regions were also evident in rare germline CG>TG SNPs from the gnomAD database^15^, indicating this phenomenon is not restricted to cancer **(Extended Data Fig. 5c)**.

**Figure 4:**
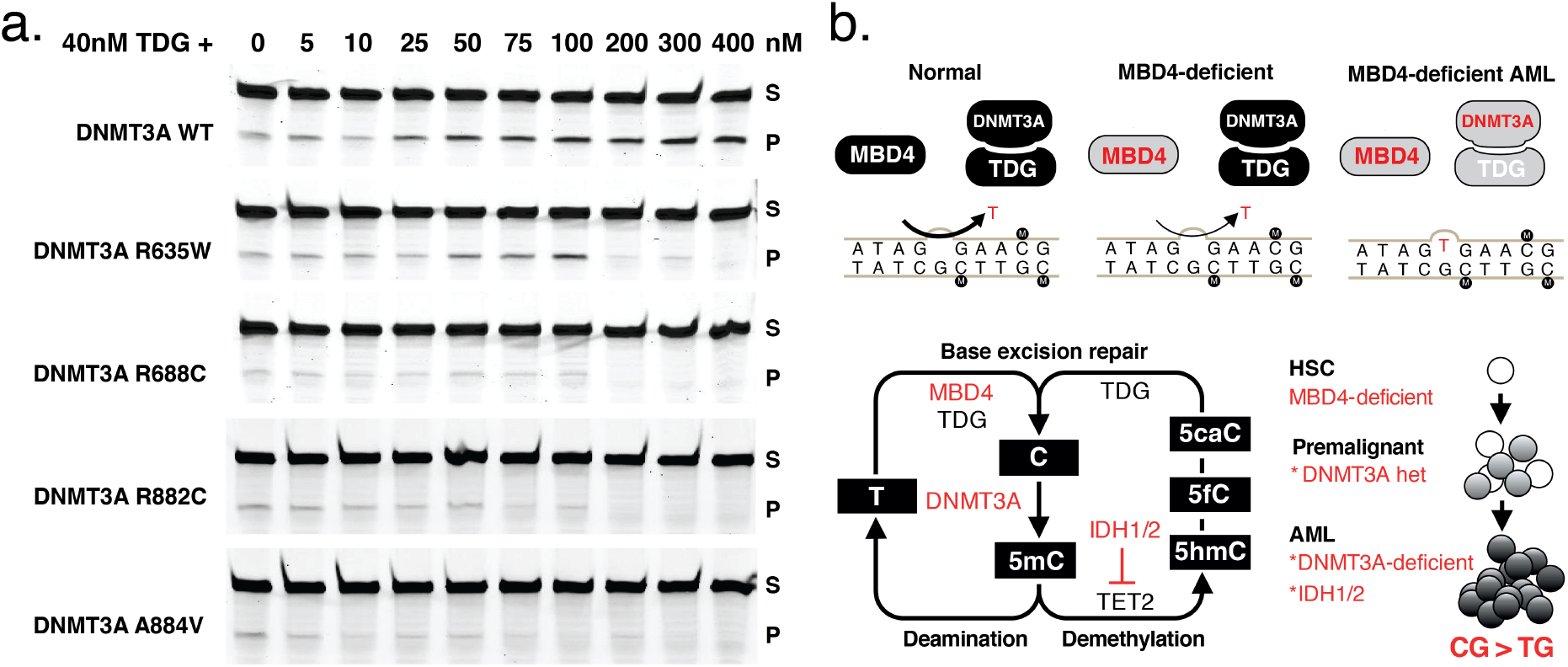
DNMT3A contributes to BER through interaction with TDG. **a**, Assessment of TDG glycosylase (40nM) activity in the presence of wildtype and mutant forms of DNMT3A, at increasing concentration (0-400nM). Substrate (S) and Product (P). Consistent results were obtained in 4 independent experiments. **b**, Schematic representation of the repair pathways governing T:G mismatch repair and the combined influence of germline mutations in MBD4 and somatic mutations in DNMT3A (at top) in AML. Schematic representation of the 5mC pathway, including deamination repair and active demethylation (at bottom). Genes mutated in the three MBD4-deficient AMLs are highlighted. A model is shown (at right), to convey the dual role of DNMT3A in driving clonal expansion and genomic instability.

The three cases with germline MBD4-deficiency shared a common path to AML. They acquired biallelic *DNMT3A* mutations and *IDH1*/*IDH2* hotspot mutations, all of which were CG>TG **(Fig. 3)**. Biallelic *DNMT3A* mutations are uncommon in AML, affecting ~3% of patients in TCGA-AML, and when considering they also have coincident *IDH1/IDH2* mutations, it is highly unlikely that these three individuals share this pattern of driver mutations by chance. These mutations impact 5mC at multiple levels – deposition (DNMT3A), removal (IDH1/IDH2) and repair (MBD4) – and this convergence suggests that modulating DNA methylation is central to AML pathogenesis in MBD4-deficient cases. Analysis of sequential bone marrow biopsies taken during treatment and single cell genotyping allowed us to refine the order of somatic mutation acquisition in two cases (EMC-AML-1, WEHI-AML-1) **(Fig. 3a-b, Extended Data Fig. 6)**. *DNMT3A* mutations present in the AML at diagnosis were also detected in non-malignant bone marrow populations in both cases, indicating that these mutations are among the first acquired. Mutations in *DNMT3A* enhance the self-renewal capacity of haematopoietic stem cells (HSCs) and are associated with age-related clonal haematopoiesis^16^–^19^. In both patients, marked expansion of clones carrying *DNMT3A* mutations occurred with treatment **(Fig. 3a-b)**, suggesting a strong advantage over normal HSCs. EMC-AML-1 experienced multiple clonal outgrowths, with nine distinct *DNMT3A* mutations, and repeated selection of clones with biallelic mutations. This shift in functional activity – the expansion of *DNMT3A*-mutant clones – increases the likelihood that cells with biallelic *DNMT3A* mutations will emerge, which appears to be key for initiating AML in MBD4-deficient patients.

There is a marked discrepancy between the substantial mutation burden in MBD4-deficient AMLs and the modest 2-3 fold increase in mutation rate in MBD4-deficient mice^20^,^21^. It is unclear whether this difference is a reflection of longer disease latency in humans, as compared to mice, or whether somatic mutations in the AML further compromise DNA repair. Mutations in *DNMT3A* and *IDH1/IDH2* have been associated with altered DNA repair in model systems^22^,^23^. It also remains unclear why TDG, a glycosylase with overlapping substrate specificity, does not compensate for MBD4 loss. One possible explanation stems from the observation that DNMT3A/B can directly stimulate TDG glycosylase activity^24^,^25^. We confirmed that recombinant DNMT3A enhances TDG glycosylase activity *in vitro* **(Fig. 4a),** but had no impact on MBD4 glycosylase activity **(Extended Data Fig. 7)**. Mutant forms of DNMT3A showed weaker stimulation, and even inhibit TDG at higher concentrations **(Fig. 4a)**. We propose a model for AML pathogenesis whereby inhibition of DNMT3A contributes in two ways: loss of one allele enables expansion of a premalignant clone, then acquisition of a second *DNMT3A* mutation increases the CG>TG mutation rate due to impaired TDG activity **(Fig. 4b)**. Supporting this model, the premalignant clone identified in WEHI-AML-1, which had a monoallelic *DNMT3A* mutation, did not carry additional mutations that would suggest an elevated mutation rate. The sporadic cancers that became MBD4-deficient (TCGA-UVM-1 and TCGA-GBM-1) did not acquire mutations in *DNMT3A* or *IDH1/IDH2,* which may indicate that this interaction is specific to the haematopoietic compartment.

The last five years have seen a concerted effort to define mutational processes that shape the cancer genome^5^. Deamination of 5mC is the most common source of somatic mutations and this damage continues to accumulate with age^26^. Our results highlight the important role for MBD4 in safeguarding against damage wrought by 5mC deamination. One manifestation of this damage is clonal haematopoiesis, a phenomenon typically observed in people >70 years of age. Individuals with biallelic loss of *MBD4* in the germline sustain high levels of damage from 5mC deamination and experience clonal expansions decades earlier, which eventually progress to AML. There are more than 40 million 5mC residues in the genome, yet these individuals develop the same type of cancer – AML – with a common set of driver mutations. Our results indicate this convergence results from the combination of a highly restricted mutational signature, which accesses a select set of driver genes, and the dual role of DNMT3A, which regulates HSC function and directly contributes to DNA repair. This interaction between mutational process, driver landscape and stem cell biology has broad implications, and may explain the tissue restricted pattern of disease in this and other cancer predisposition syndromes.

## Acknowledgements

The authors would like to thank Simon He, Anita Rijneveld, Kirsten van Lom and Kirsten Gussinklo for providing clinical information and reviewing samples; Meaghan Wall for assistance with cytogenetics; Naomi Sprigg for assistance with sample collection; Elwin Rombouts for assistance with single cell sorting; Hideharu Hashimoto and Xiaodong Cheng for the TDG expression vector; Sari van Rossum and Joyce Lebbink for assistance with recombinant protein isolation; the Australasian Leukaemia and Lymphoma Group for access to clinical samples; and Stephen Wilcox for technical assistance with sequencing. Additional sequencing was performed at The Australian Genome Research Facility (Melbourne, Australia) and the Kinghorn Centre for Clinical Genomics (Sydney, Australia). Sean Grimmond, Jason Wong, Oliver Sieber, Alicia Oshlack and Stephen Nutt provided valuable feedback on the manuscript.

This work was made possible through support from the Australian National Health and Medical Research Council (NHMRC) (Program Grant 1113577, to W.S.A and A.W.R), an Independent Research Institutes Infrastructure Support Scheme Grant (9000220), a Victorian State Government Operational Infrastructure Support Grant, the Netherlands Organisation for Scientific Research (NWO) and the Center for Translational Molecular Medicine (CTMM). M.A.S is supported by a grant from CTMM (GR03O-102) and a Rubicon fellowship from NWO (019.153LW.038), E.C. is a recipient of a PhD scholarship from the Leukaemia Foundation of Australia, A.H. is a recipient of a PhD scholarship from the Ministry of Health - Sultanate of Oman, M.E.B is supported by the Bellberry-Viertel fellowship, W.S.A and A.W.R are supported by fellowships from NHMRC (1058344 and 1079560, respectively), and I.J.M. is supported by the Victorian Cancer Agency. We wish to acknowledge the generous philanthropic support of the Felton Bequest, Malcolm Broomhead, and BHP Billiton.

The results are in part based upon data generated by the TCGA Research Network (http://cancergenome.nih.gov/) and the Epigenetic studies in Acute Myeloid Leukemia [phs001027], which was supported by K08CA169055 (F.E. Garrett-Bakelman), Starr Cancer Consortium I4-A442 (A.M. Melnick, R. Levine and C.E. Mason) and LLS SCOR 7006-13 (A.M. Melnick).

## Contributions

M.A.S., E.C., A.W.R., P.J.M.V. and I.J.M. initiated the project. M.A.S. and C.F. coordinated the sequencing analysis. Processing of clinical material and preparation of sequencing libraries were performed by E.C. (WEHI-AML) and by A.Z., A.H., and F.G.K. (EMC-AML). B.Lu. and F.G.K. sorted single cells and performed WGA and genotyping by Sanger sequencing. A.Z., A.H., M.R. and F.G.K. prepared sequencing libraries from single cells for EMC-AML-1. E.M.J.B. coordinated sequencing experiments (EMC-AML). A.Z. and A.H. isolated the recombinant proteins and performed DNA glycosylase assays. E.C. and S.E.M. validated germline and somatic variants and performed phasing. C.F. and I.J.M. performed the analysis of TCGA samples. T.M. prepared RRBS libraries. Methylation data was analysed by C.F. and M.A.S. with input from E.C., M.E.B. and I.J.M. M.A.S. developed the testing framework to assess mutation rate. M.A.S. and I.J.M. refined the base context analysis. M.A.S., E.C., C.F., A.W.R., P.J.M.V. and I.J.M. wrote the manuscript with input from the other authors. W.S.A. and B.Lo. reviewed results and provided critical feedback. All authors discussed the results and agree with the conclusions presented.

## Extended Data

Germline loss of *MBD4* predisposes to leukaemia due to a mutagenic cascade driven by 5mC

**Extended Data Tables – pg. 2-3**

Extended Data Table 1 - Candidate germline loss-of-function variants in *MBD4*

Extended Data Table 2 - Clinical sample description

**Extended Data Figures – pg. 4-11**

Fig. 1 - Mutational signatures in cancers with germline loss-of-function variants in *MBD4*

Fig. 2 - Confirmation and phasing of *MBD4* variants in WEHI-AML cases

Fig. 3 - Copy number and RNA splicing in cancers with germline loss-of-function variants in *MBD4*

Fig. 4 - Methylation status at somatic mutation positions in glioblastoma

Fig. 5 - Assessment of genetic and epigenetic factors that impact mutation rate

Fig. 6 - Single cell genotyping for EMC-AML-1 at relapse

Fig. 7 - MBD4 glycosylase activity in the presence of DNMT3A

Fig. 8 - Comparison between TCGA consensus calls and superFreq output

**Clinical synopsis – pg. 12**

**Methods – pg. 13-20**

**Extended Data References – pg. 20-21**

## Supplementary Information

Somatic mutations detected in MBD4-deficient AML at diagnosis (hg19). VCFv4.0 format. A quality score is provided (SOMATICP), variants with a score >0.5 were used for mutation signature analysis. WEHI-AML-1.vcf [Diagnosis, Bone Marrow, Exome]

WEHI-AML-1.G.vcf [Diagnosis, Bone Marrow, Genome]

WEHI-AML-2.vcf [Diagnosis, Bone Marrow, Exome]

WEHI-AML-2.G.vcf [Diagnosis, Bone Marrow, Genome]

EMC-AML-1.vcf [Diagnosis, Bone Marrow, Exome]

Average FPKM values from WEHI-AML-1 and WEHI-AML-2. Columns are chromosome, start, end, gene, average FPKM, strand.

WEHI-AML-FPKM-expression.bed

Calculation of relative mutation rates for WEHI-AML-1 and WEHI-AML-2. This analysis only includes mutations covered in the WGBS controls. Supplementary_Information_RMR.xlsx

**Extended Data Table 1.** Candidate germline loss-of-function variants in *MBD4*. Code: Sample identifiers, TCGA samples were re-coded. Genomic position in hg19 (chr: Chromosome. pos: Position. ref: reference. var: variant. dbSNP: rs identifier). Coverage values derived from exome data for the cancers and matched normal samples. An estimate of tumour purity was calculated based on copy number alterations and somatic SNV frequency. Annotation: Gene, transcript, annotation and variant type.

| code | cancer type | chr | pos | ref | var | normal<br>ref var | cancer<br>ref var | tumour<br>content | dbSNP | gene | transcript | annotation | type |
| --- | --- | --- | --- | --- | --- | --- | --- | --- | --- | --- | --- | --- | --- |
| EMC-AML-1 | Acute Myeloid Leukaemia | 3 | 129150385 | ATG | - | 1 11 | 2 12 | 90% | rs775848563 | MBD4 | NM_003925 | p.His567del | inframe_deletion |
| SW1783 | Glioma (cell line) | 3 | 129150399 | A | T |  |  |  | rs200758755 | MBD4 | NM_003925 | p.Leu563Ter | stop_gained |
| TCGA-GBM-2 | Glioblastoma Multiforme | 3 | 129150399 | A | T | 105 80 | 127 108 | 90% | rs200758755 | MBD4 | NM_003925 | p.Leu563Ter | stop_gained |
| TCGA-OV-2 | Ovarian Serous Cystadenocarcinoma | 3 | 129150399 | A | T | 74 53 | 48 232 | 90% | rs200758755 | MBD4 | NM_003925 | p.Leu563Ter | stop_gained |
| TCGA-UCEC-1 | Uterine Corpus Endometrial Carcinoma | 3 | 129151375 | G | A | 19 26 | 26 57 | 90% |  | MBD4 | NM_003925 | p.Arg546Ter | stop_gained |
| TCGA-OV-1 | Serous Ovarian Cancer | 3 | 129151450 | C | A | 70 74 | 109 215 | 90% | rs778697654 | MBD4 | NM_003925 | c.1562-1G>T | splice_acceptor |
| WEHI-AML-1 | Acute Myeloid Leukaemia | 3 | 129151450 | C | A | 26 25 | 41 32 | 80% | rs778697654 | MBD4 | NM_003925 | c.1562-1G>T | splice_acceptor |
| WEHI-AML-2 | Acute Myeloid Leukaemia | 3 | 129151450 | C | A | 42 28 | 24 38 | 80% | rs778697654 | MBD4 | NM_003925 | c.1562-1G>T | splice_acceptor |
| TCGA-UVM-1 | Uveal Melanoma | 3 | 129151450 | C | A | 43 43 | 1 32 | 90% | rs778697654 | MBD4 | NM_003925 | c.1562-1G>T | splice_acceptor |
| TCGA-THYM-1 | Thymoma | 3 | 129152813 | G | A | 14 28 | 35 31 | <10% |  | MBD4 | NM_003925 | p.Arg431Ter | stop_gained |
| WEHI-AML-1 | Acute Myeloid Leukaemia | 3 | 129155547 | C | CT | 84 45 | 133 64 | 80% | rs558765093 | MBD4 | NM_003925 | p.Val314ArgfsTer13 | frameshift |
| WEHI-AML-2 | Acute Myeloid Leukaemia | 3 | 129155547 | C | CT | 126 84 | 117 69 | 80% | rs558765093 | MBD4 | NM_003925 | p.Val314ArgfsTer13 | frameshift |
| TCGA-STAD-1 | Stomach adenocarcinoma | 3 | 129155723 | G | C | 39 38 | 98 85 | 70% | rs749739092 | MBD4 | NM_003925 | p.Ser255Ter | stop_gained |
| TCGA-GBM-1 | Glioblastoma Multiforme | 3 | 129156562 | C | T | 51 67 | 15 55 | 90% | rs552296498 | MBD4 | NM_003925 | c.335+1G>A | splice_donor |
| TCGA-CHOL-1 | Cholangiocarcinoma | 3 | 129156562 | C | T | 13 15 | 31 22 | 80% | rs552296498 | MBD4 | NM_003925 | c.335+1G>A | splice_donor |

**Extended Data Table 2.** Clinical sample description. Description of clinical samples and molecular profiling approaches applied to each. HSCT: Haematopoietic stem cell transplant, BM: Bone marrow, PB: Peripheral blood, AML: Acute myeloid leukaemia, MDS: Myelodysplastic syndrome, WGS: Whole genome sequencing, WES: Whole exome sequencing, RNA-seq: RNA sequencing, RRBS: Reduced representation bisulfite sequencing, eRRBS: Enhanced reduced representation bisulfite sequencing, Panel: Illumina TruSight Myeloid Sequencing Panel.

| Sample | Time from diagnosis days (months) | Clinical time point | Tissue | Blast % morphology | Pathology summary | Analysis |
| --- | --- | --- | --- | --- | --- | --- |
| 1 | 0 (0) | Diagnosis | BM | 21 | Diagnosis of AML | WGS, WES, RNA-seq, RRBS |
| 2 | 35 (1) | Post induction | BM | 3 | In morphologic and cytogenetic remission | WGS, WES, RNA-seq, RRBS |
| 3 | 69 (2) | Post consolidation cycle 1 | BM | 2 | In morphologic and cytogenetic remission | WES, RNA-seq, RRBS |
| 4 | 115 (3) | Post consolidation cycle 2 | BM | 4 | In morphologic and cytogenetic remission | WES, RNA-seq |
| 5 | 148 (4) | Pre allogeneic HSCT | BM | 13 | Relapsed AML | WES, RNA-seq |
| 6 | 202 (6) | Post allogeneic HSCT | BM | 3.5 | In morphologic and cytogenetic remission | WES |
| 7 | 245 (8) | Relapse | BM | 36 | Relapsed AML | WES, RNA-seq, RRBS |

| Sample | Time from diagnosis days (months) | Clinical time point | Tissue | Blast % morphology | Pathology summary | Analysis |
| --- | --- | --- | --- | --- | --- | --- |
| 1 | -1410 (-46) | HSCT donor to WEHI-AML-1 | PB | 0 | Normal full blood count and blood film | WES |
| 2 | 0 | Diagnosis | BM | 22 | Diagnosis of AML | WGS, WES, RNA-seq, RRBS |
| 3 | 33 (1) | Post induction | BM | 1.5 | In morphologic and cytogenetic remission | WGS, WES, RNA-seq, RRBS |

**EMC-AML-1 with AML, not otherwise specified (acute monocytic leukaemia, WHO ICD 9891/3)**
| Sample | Time from diagnosis days (months) | Clinical time point | Tissue | Blast % morphology | Pathology summary | Analysis |
| --- | --- | --- | --- | --- | --- | --- |
| 1 | 0 | Diagnosis | BM | 84 | Diagnosis of AML | WES, RNA-Seq, Panel, eRRBS |
| 2 | 27 (1) | Induction cycle 1 (Day 25) | BM | 32 | Persistent AML | Panel |
| 3 | 48 (1) | Induction cycle 2 (Day 17) | BM | 3 | Hypocellular bone marrow | Panel |
| 4 | 55 (2) | Induction cycle 2 (Day 25) | BM | 0.8 | In morphologic remission | Panel |
| 5 | 64 (2) | Complete remission | BM | 0.4 | In morphologic remission | Panel |
| 6 | 77 (3) | Pre autologous HSCT | BM | 0.8 | In morphologic remission | Panel |
| 7 | 125 (4) | Post autologous HSCT | BM | 0.6 | In morphologic remission | Panel |
| 8 | 169 (6) | Complete remission | BM | 1.2 | In morphologic remission | Panel |
| 9 | 253 (8) | Complete remission | BM | 1.8 | In morphologic remission | Panel |
| 10 | 351 (12) | Complete remission | BM | 0.4 | In morphologic remission | Panel |
| 11 | 496 (16) | Complete remission | BM | 1.6 | In morphologic remission | Panel |
| 12 | 841 (28) | Relapse timepoint 1 | BM | 16.2 | Relapsed AML | Panel, Single cells |
| 13 | 853 (28) | Relapse timepoint 2 | BM | 7.6 | Relapsed AML | WES, RNA-Seq, Panel |
| 14 | 877 (29) | Complete remission | BM | 0.8 | In morphologic remission | Panel |
| 15 | 890 (29) | Complete remission | BM | 2.4 | In morphologic remission | Panel |

**Extended Data Figure 1.**
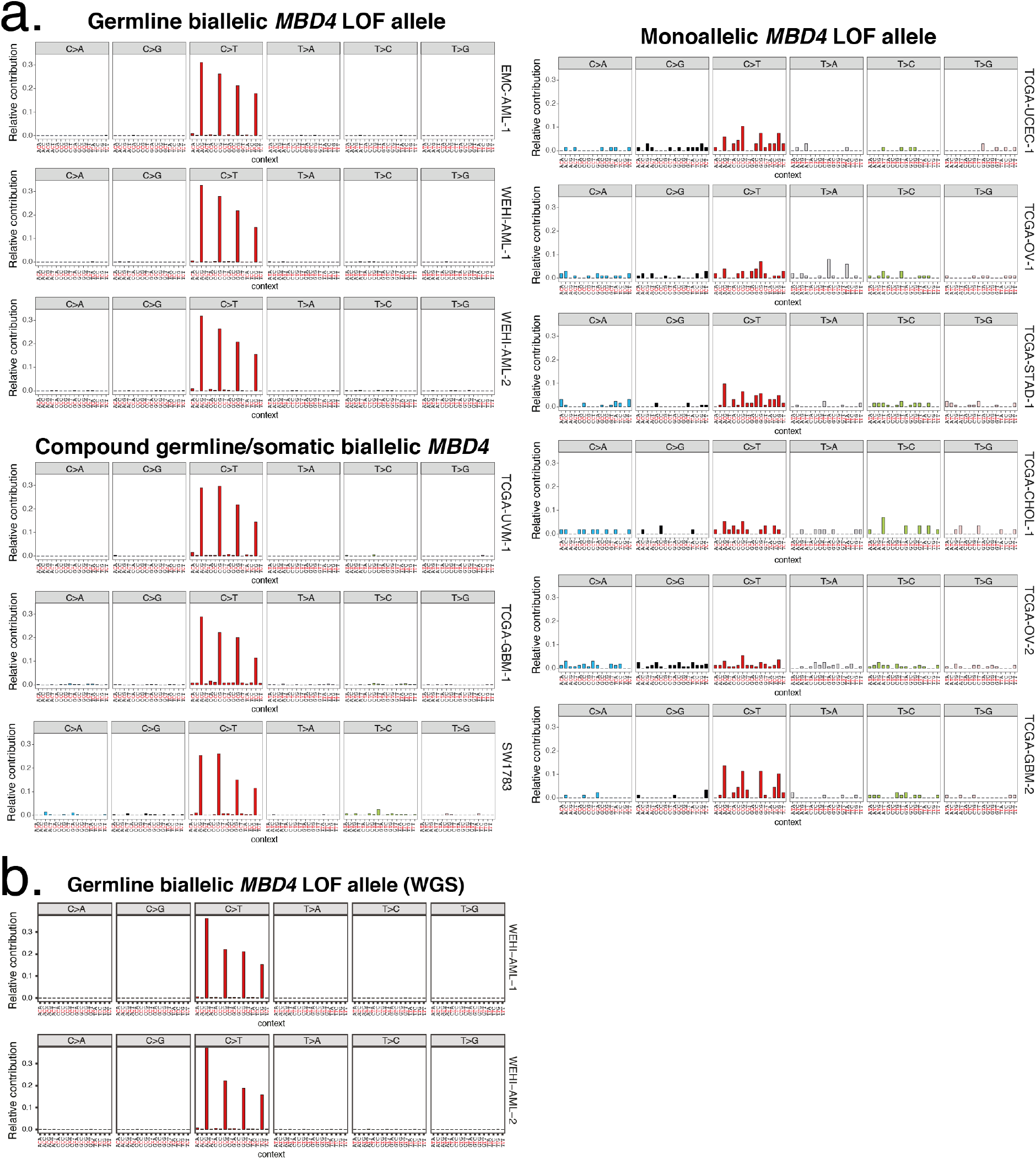
Mutational signatures in cancers with germline loss-of-function variants in *MBD4*. **a**, Trimer mutation profiles from exome data from the diagnostic AMLs (EMC-AML-1, WEHI-AML-1 and WEHI-AML-2), cancers from TCGA with germline loss-of-function variants and the SW1783 cell line (COSMIC). Samples with biallelic *MBD4* inactivation are shown at left (separated into germline biallelic and compound germline/somatic), and cases with monoallelic *MBD4* inactivation are shown at right. TCGA-THYM-1 was excluded due to low purity. There was a significant enrichment of mutations in the ACG context compared to TCG in MBD4-deficient exomes (p=0.007937, Mann-Whitney U test, Supplementary Information). **b**, Trimer context for somatic mutations identified in whole genome sequencing (WGS) from WEHI-AML-1 (n=14,313 mutations) and WEHI-AML-2 (n=15,543 mutations).

**Extended Data Figure 2.**
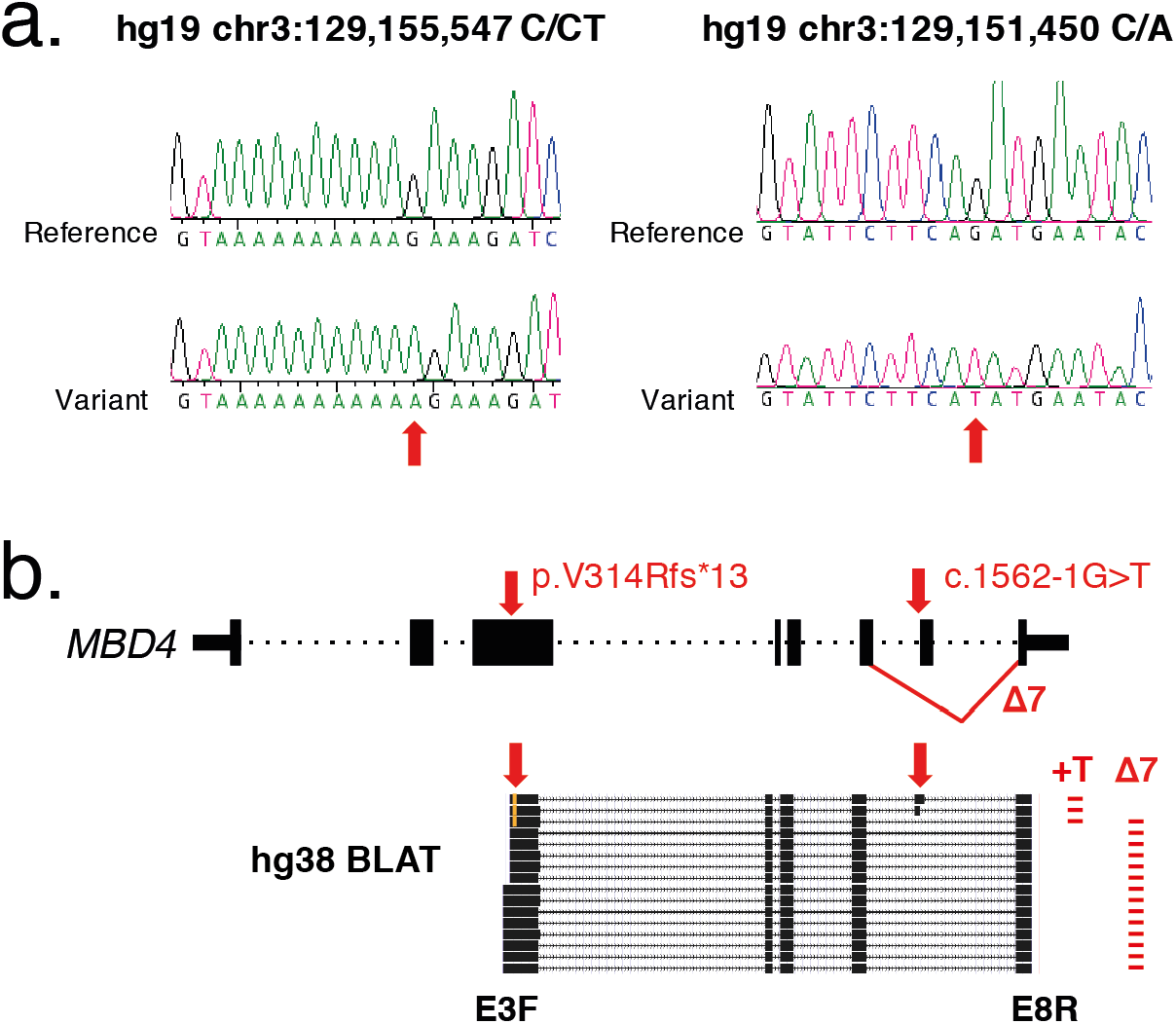
Confirmation and phasing of *MBD4* variants in WEHI-AML cases. **a**, Confirmation of the *MBD4* variants in WEHI-AML-1 and WEHI-AML-2. Sanger sequencing traces were generated from cloned PCR products after amplification from DNA (top). The reverse complement of the genome sequence is shown, which represents the open reading frame 5’-3’. **b**, A schematic of the *MBD4* gene is shown. PCR was performed on cDNA to generate fragments that covered both variant positions, using primers in exon 3 and exon 8. Products cloned into TOPO were sequenced and BLAT results are shown to demonstrate transcript structure. Each product was assessed for the presence of the +T insertion (+T) and aberrant splicing of exon 7 *(Δ7).* The variants were mutually exclusive in fifteen out of sixteen clones. In one clone the variants were coincident, likely due to instability in the poly-T stretch (during reverse transcription or subsequent PCR amplification).

**Extended Data Figure 3.**
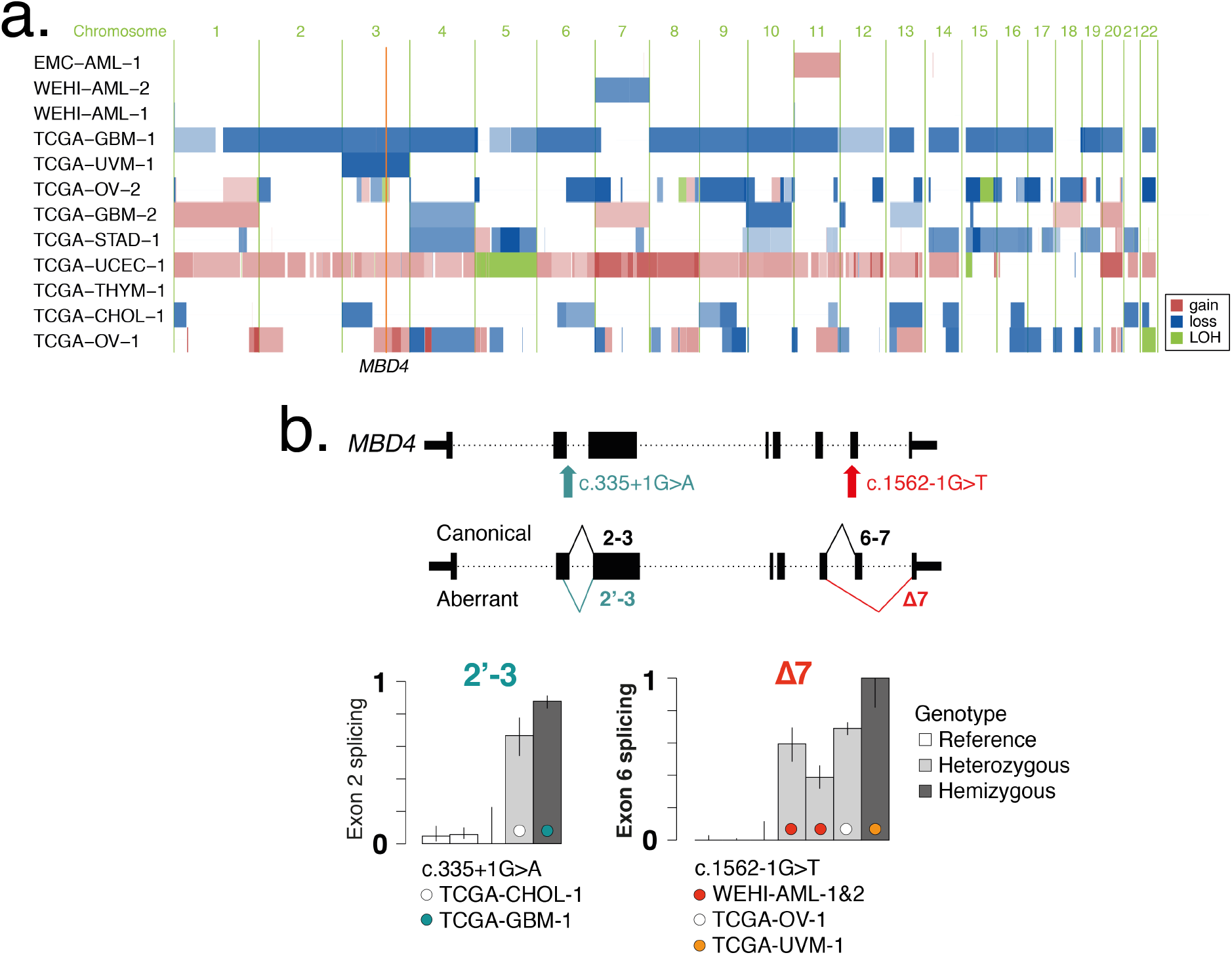
Copy number and RNA splicing in cancers with germline loss-of-function variants in *MBD4*. **a**, Copy number profiles were generated from cancer exome data by tracking shifts in allele frequencies at heterozygous germline SNPs and coverage (summarised at the gene level). Sample codes are shown, gain is represented in red, loss in blue, and loss of heterozygosity in green. The position of *MBD4* is highlighted. The intensity of the colour is proportional to the clonality, with darker colours representing higher clonality estimates. **b**, A schematic of the *MBD4* gene is shown at top together with the position of two candidate loss-of-function variants that impact splice sites. The genotype status is indicated, which is informed based on the copy number calls. Canonical and aberrant splice products are shown below, which were quantified with RNA-seq by counting reads that span splice junctions. Three unrelated control samples were selected to serve as controls for canonical splicing. Each bar represents data from a single sample. The error bars reflect the level of coverage and are set at 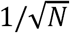, where *N* is the total number of spliced reads.

**Extended Data Figure 4.**
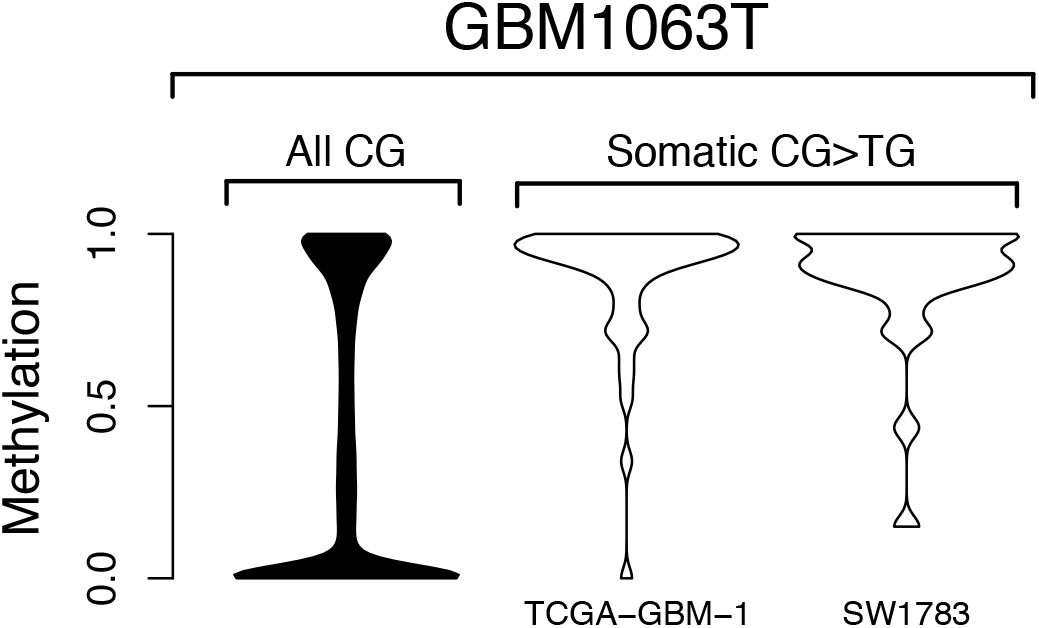
Methylation status at somatic mutation positions in glioblastoma. Methylation status was extracted for CG sites from RRBS data from an unrelated glioblastoma (GBM1063T)^1^, either at all CG sites (All CG) or at sites of somatic CG>TG mutations in TCGA-GBM-1 or the cell line SW1783. Sites with mutations were typically fully methylated in the control sample.

**Extended Data Figure 5.**
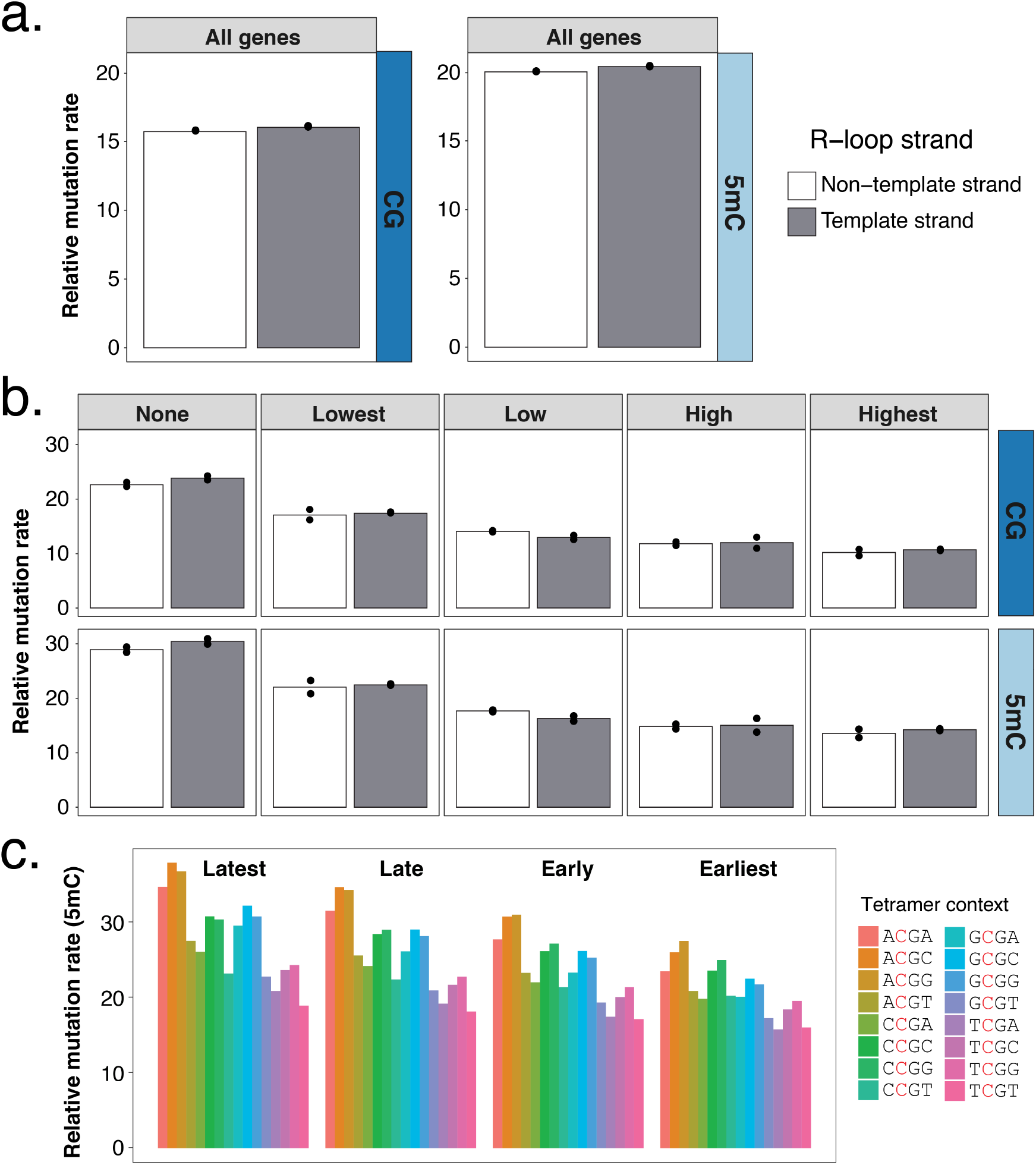
Assessment of genetic and epigenetic factors that impact mutation rate. **a**, Mutations from WEHI-AML-1 and WEHI-AML-2 were tested for association with transcriptional strand. The relative mutation rate was calculated per megabase of CG sites (dark blue) or per megabase of 5mCG sites (light blue). The values are corrected to account for the total mutation load (see Methods). Individual values are plotted (n=2) and the bar shows the mean. **b**, Genes were divided into bins based on their expression (see Methods). The relative mutation rate was calculated per bin based on CG or 5mCG abundance (as in **a**). Individual values are plotted (n=2) and the bar shows the mean. There is considerable correlation between replication domain and expression level. **c**, The NCGN tetramer context was determined for rare germline CG>TG SNPs from the gnomAD database^2^. The dataset was split based on replication timing, with a quarter of the genome ascribed to each bin. Note that the correction for methylation status uses data derived from CD34+ blood cells.

**Extended Data Figure 6.**
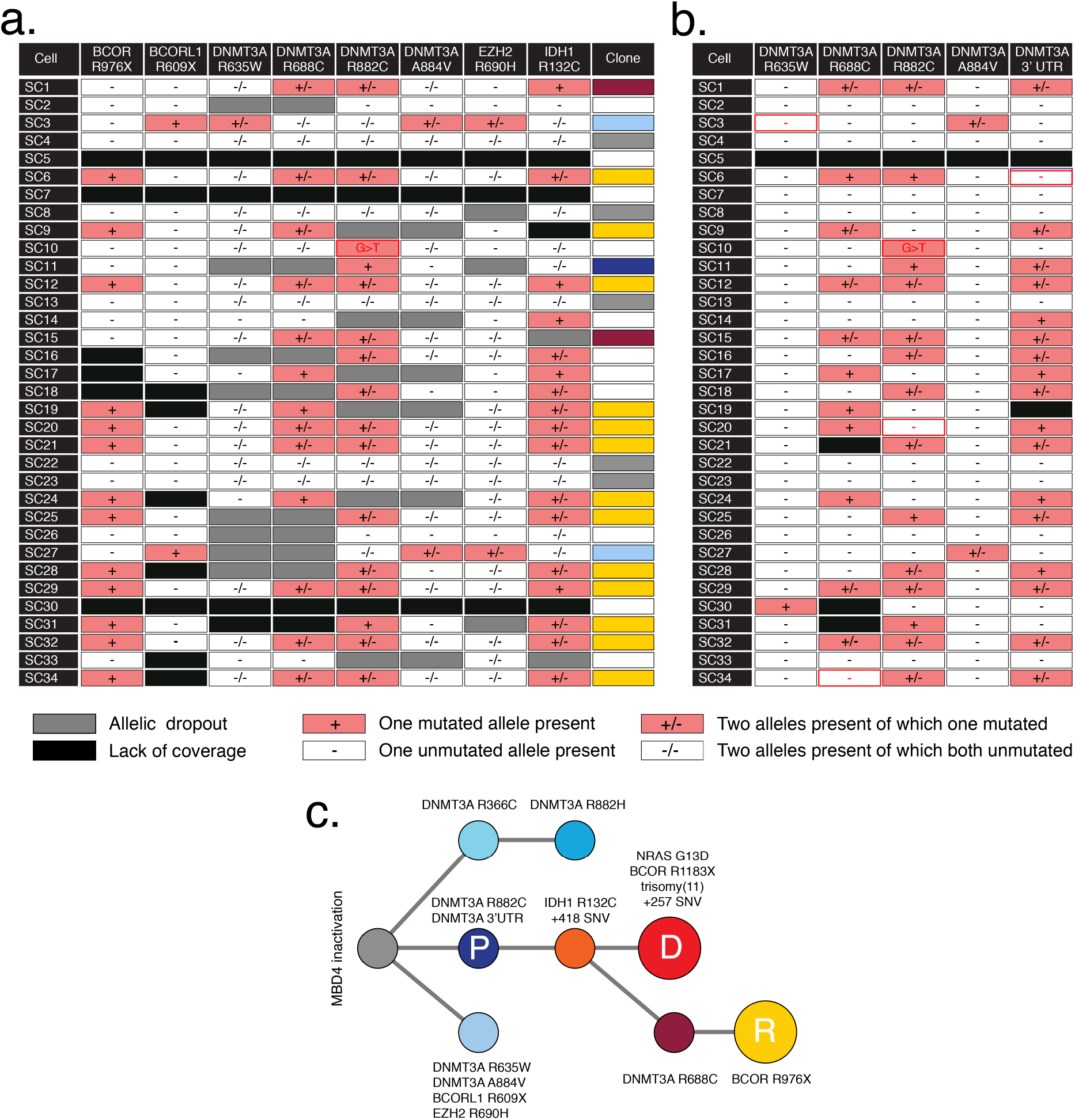
Single cell genotyping for EMC-AML-1 at relapse. Single cells isolated at relapse (SC1-SC34, Sample #12, 841 days post-diagnosis) were genotyped for key mutations after whole genome amplification. Cells were assessed using the TruSight Myeloid Sequencing Panel (**a**), or by PCR followed by Sanger sequencing (**b**). The genotyping results were used to assigned each cell to a clone, with colours matching the phylogenetic tree presented in Figure 3 (repeated here for clarity (**c**). No clonal assignment was made if the genotyping results were ambiguous due to technical failure (white). Discordant results between the platforms are highlighted using red text and outline (in **b**) and generally reflect the lower sensitivity of detection for Sanger sequencing. For SC10 multiple alleles were detected at the site of the DNMT3A R882C mutation (denoted with a G>T symbol).

**Extended Data Figure 7.**
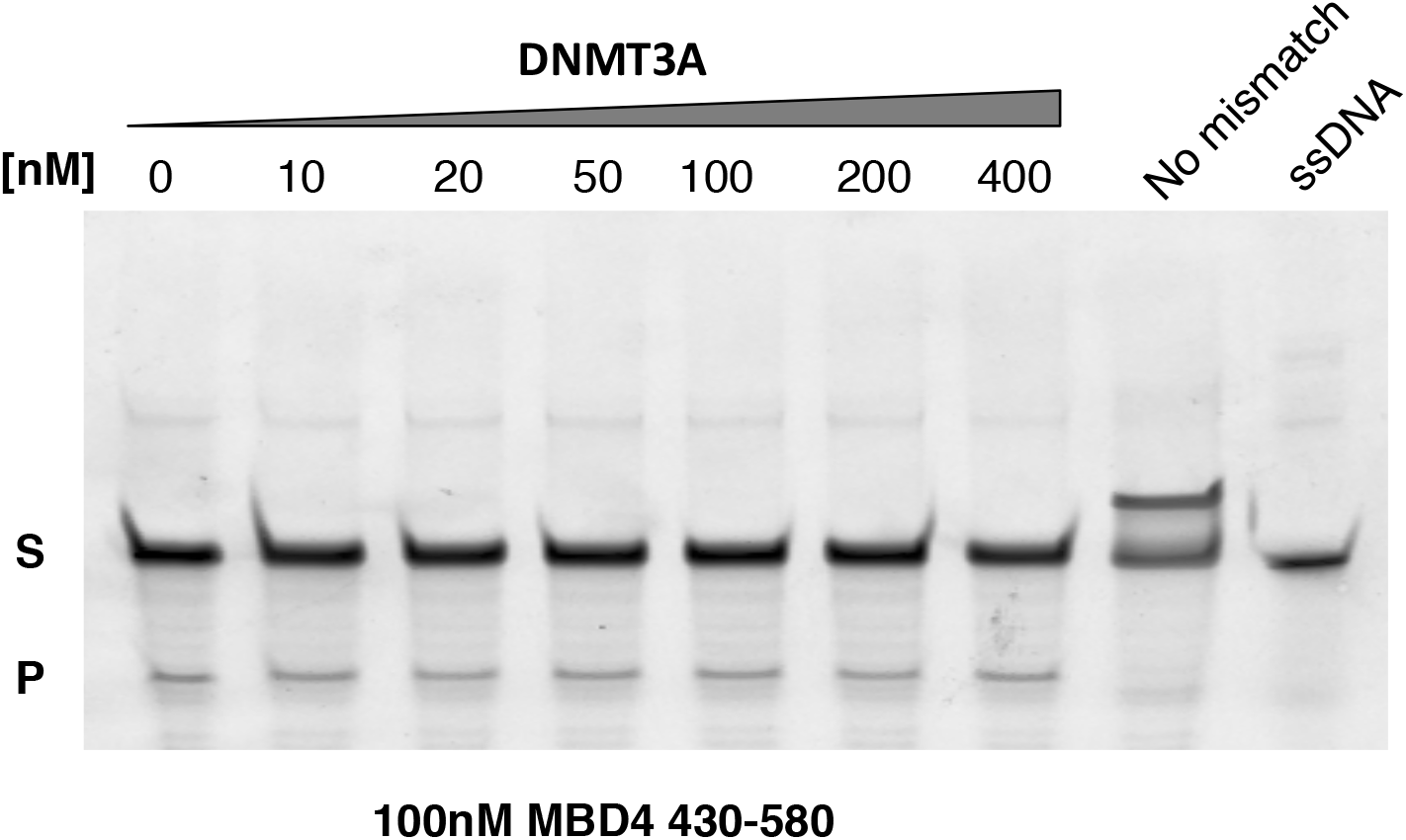
MBD4 glycosylase activity in the presence of DNMT3A. A glycosylase assay was performed to assess the activity of MBD4 glycosylase domain (AA430-580) in the presence of increasing amounts of wildtype DNMT3A (0-400nM). A template without a mismatch (No mismatch) and single stranded DNA were included as negative controls, as they should not be cleaved. Substrate (S) and Product (P). Consistent results were obtained in 2 independent experiments.

**Extended Data Figure 8.**
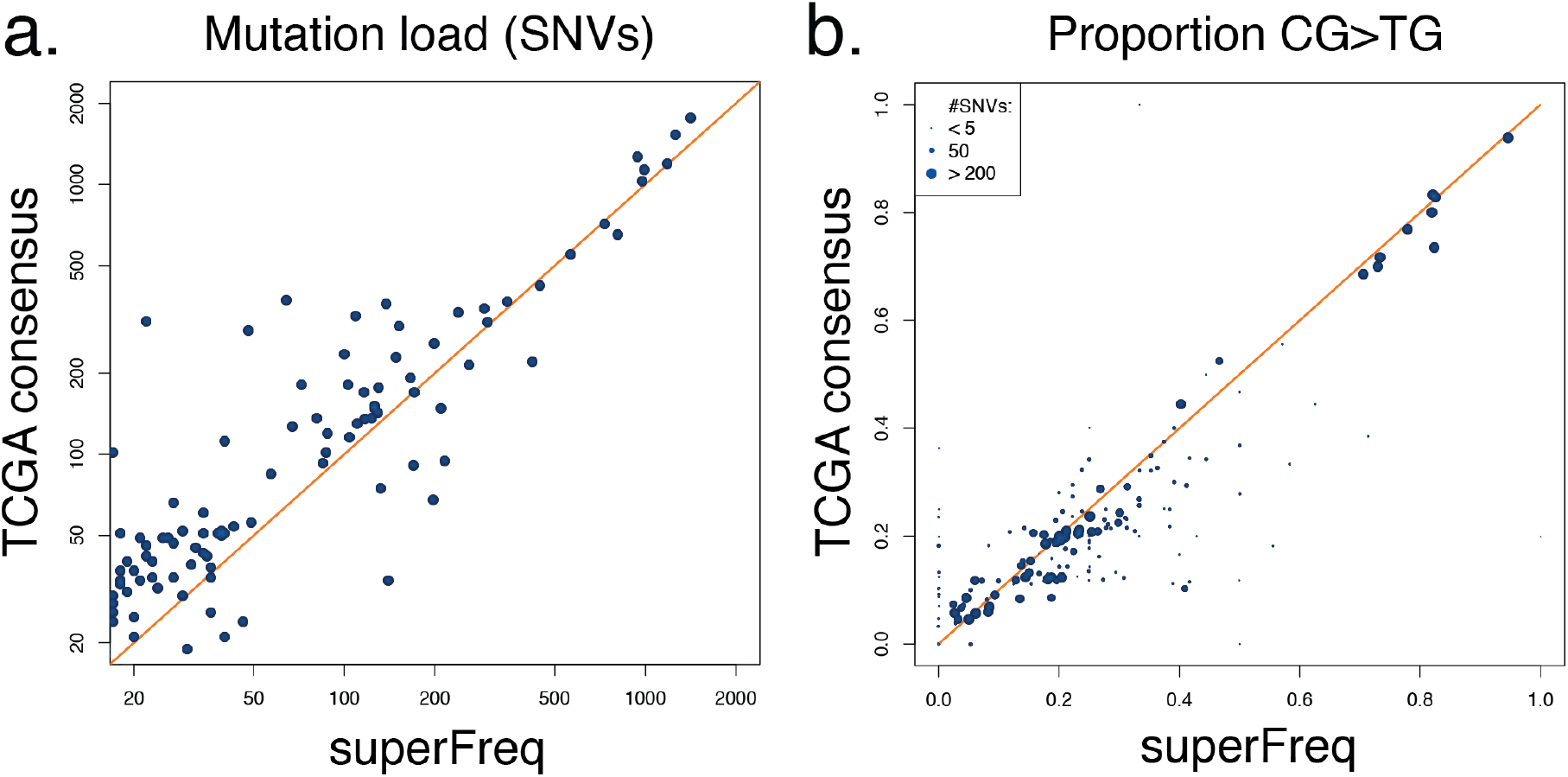
Comparison between TCGA consensus calls and superFreq output. In Figure 1D we included estimates of mutation load and proportion of CG>TG mutations from two sources. The values for WEHI-AML and EMC-AML samples were generated using our own analysis platform, superFreq, whereas the data from the 10,683 TGCA cases was generated from consensus calls available from GDC (see Methods). To ensure the results were comparable, we used superFreq to analyse a subset of TCGA samples, which included all of the TCGA-AML cases and the TCGA samples with the highest proportion of CG>TG mutations based on the consensus calls. We observed a satisfactory correlation between the results generated using superFreq and those from the TCGA consensus calls, both for mutation load (**a**) and CG>TG proportion (**b**). In **b** the point size is scaled to reflect the total number of mutations.

## Clinical synopsis

EMC-AML-1 was 33 years old when diagnosed with acute monocytic leukaemia (AML, WHO ICD 9891/3). The AML had trisomy 11 on karyotyping and was negative for *NPM1, FLT3* and *CEBPα* mutations. His medical history included colonic polyps requiring a hemicolectomy 2 years prior to his AML diagnosis. His AML was refractory to induction chemotherapy (standard dose cytarabine and daunorubicin). Repeat induction with intermediate dose cytarabine resulted in complete morphologic and cytogenetic remission. He then had an autologous haematopoietic stem cell transplant (HSCT) with BU-CY conditioning (busulfan and cyclophosphamide). He relapsed 2 years and 3 months post autologous HSCT. The AML at relapse had a normal karyotype and was negative for *NPM1, FLT3* and *CEBPα* mutations. Salvage induction chemotherapy (high dose cytarabine, mitoxantrone and etoposide) resulted in complete morphologic remission. This was followed by an allogeneic HSCT from a matched unrelated donor with myeloablative and total body irradiation conditioning. He achieved complete morphologic remission with full donor chimerism. He developed extensive graft-versus-host disease (GVHD) with secondary graft failure responsive to steroids and EBV-reactivation requiring rituximab. He died 2 years post allogeneic HSCT with relapsed AML.

WEHI-AML-1 was 31 years old when diagnosed with AML with myelodysplasia-related changes (MDS-associated cytogenetic abnormality, monosomy 7, WHO ICD 9895/3). The AML was negative for *NPM1, FLT3* and *CEBPa* mutations. She had induction chemotherapy (high dose cytarabine, idarubicin and etoposide) and achieved complete morphologic and cytogenetic remission. This was followed by 2 cycles of consolidation chemotherapy (standard dose cytarabine, idarubicin and etoposide). Early morphologic relapse was detected on bone marrow examination prior to allogeneic HSCT from her female sibling (WEHI-AML-2) with BU-CY conditioning. Bone marrow examination 5 weeks post allogeneic HSCT showed complete morphologic and cytogenetic remission; and full donor chimerism. Relapsed AML (of WEHI-AML-1 origin) occurred 11 weeks post allogeneic HSCT. Salvage therapy with FLAG chemotherapy regimen (fludarabine, cytarabine and filgrastim) proved unsuccessful. WEHI-AML-1 died of relapsed AML less than 12 months after diagnosis.

WEHI-AML-2 was 30 years old when she donated peripheral blood stem cells to WEHI-AML-1. Her medical history included iron deficiency anaemia secondary to menorrhagia and bleeding from descending colon and rectal polyps. Her full blood count was normal at the time of stem cell donation. Her routine full blood count 4 years later, at 34 years old, showed pancytopenia. A diagnosis of AML with myelodysplasia-related changes (MDS-associated cytogenetic abnormality, monosomy 7, WHO ICD 9895/3) was made on bone marrow examination. The AML was negative for *NPM1, FLT3* and *CEBPa* mutations. She had induction chemotherapy (high-dose cytarabine and idarubicin) and achieved complete morphologic and cytogenetic remission. This was followed by one cycle of consolidation chemotherapy (standard dose cytarabine, idarubicin and etoposide). She then had an allogeneic HSCT using two partially HLA-matched umbilical cord blood units following FLU-CY-TBI conditioning (fludarabine; cyclophosphamide and total body irradiation). She developed grade 1 GVHD of the gut. She remains in complete morphologic and cytogenetic remission.

## Methods

### Patient characteristics and sample collection

EMC-AML-1, WEHI-AML-1 and WEHI-AML-2 were diagnosed with AML and treated with combination chemotherapy as per the protocols at their respective institutions **[see Clinical Synopsis].** They gave informed consent according to the Declaration of Helsinki for participation in research and for collection of samples over the course of their treatment. This research project was approved by our respective Human Research Ethics Committees (MEC-2015-155, WEHI HREC 13/01 and MH-HREC 2012.274). Samples from bone marrow and peripheral blood were collected over the course of their treatment **[Extended Data Table 2]**. In addition, WEHI-AML-2 acted as a donor for an allogeneic haematopoietic stem cell transplant for WEHI-AML-1 and had peripheral blood taken for chimerism analysis at time of donation that was available for molecular analysis.

### Whole exome sequencing and whole genome sequencing

Whole exome sequencing on EMC-AML-1 was performed as previously described^3^. For WEHI-AML-1 and WEHI-AML-2, DNA was extracted using either the MagAttract DNA Blood M48 Kit or QIAsymphony DSP DNA Mini Kit (Qiagen, Venlo, the Netherlands). DNA libraries were made using the Illumina TruSeq Nano DNA Sample Preparation Kit (Illumina, San Diego, CA, USA) as per manufacturer’s instructions. Briefly, between 50 to 100 ng of DNA was sheared using a Covaris S220 (Covaris, Woburn, MA, USA) to generate DNA insert sizes of 350bp. Indexed adapters were ligated onto the DNA fragments and the library was enriched with 8 cycles of PCR. DNA libraries were quantified and used for both whole genome sequencing and whole exome sequencing. Whole genome sequencing was performed on a HiSeq X Ten (Illumina, San Diego, CA, USA) by The Kinghorn Centre for Clinical Genomics (KCCG, Sydney, Australia). For the exome capture, a pool of eight indexed DNA libraries was hybridised with the Human All Exon v5+UTR Capture Library using the SureSelect^*XT2*^ Target Enrichment System (Agilent Technologies, Santa Clara, CA, USA). Captured libraries were amplified with 9 cycles of PCR, quantified, diluted and sequenced on a HiSeq2500 (Illumina, San Diego, CA, USA) at the Australian Genome Research Facility (AGRF) (Melbourne, Australia).

### Reduced representation bisulfite sequencing (RRBS)

For WEHI-AML-1 and WEHI-AML-2, between 75 to 100 ng of DNA was used to construct RRBS libraries using the Ovation RRBS Methyl-Seq System (NuGEN, San Carlos, CA, USA). DNA was restriction enzyme digested using Mspl followed by ligation with indexed adaptors. Bisulfite conversion was performed using the Epitect kit (Qiagen, Venlo, the Netherlands). RRBS libraries were amplified with either 12 or 13 cycles of PCR for 100 or 75 ng of initial DNA respectively. The libraries were quantified, diluted and sequenced on a HiSeq2500 at AGRF. Enhanced RRBS data from EMC-AML-1 was available through the Database of Genotypes and Phenotypes (dbGaP) (phs001027) and RRBS data from a glioblastoma, GBM1063T, was available from Gene Expression Omnibus (GSE70175)^1^. RRBS sequencing reads were first trimmed to remove adapters and low-quality sequence (base quality <20) with Trim_Galore. Diversity adaptors were removed with a python script provided by NuGEN, (trimRRBSdiversityAdaptCustomers.py). Alignment to hg19 was performed with Bismark 0.13.0 and methylation status was assessed using bismark_methylation_extractor, ignoring the 5 bases at the 5’ end of each read^4^.

### RNA sequencing

For WEHI-AML-1 and WEHI-AML-2, total RNA was extracted using TRIzol (Thermo Fisher Scientific, Waltham, MA, USA) as per manufacturer’s instructions. RNA libraries were generated using the TruSeq RNA Sample Preparation Kit v2 (Illumina, San Diego, CA, USA). The libraries were made using 1000ng of total RNA as input, except for the WEHI-AML-1 diagnosis time point, where only 170ng was available. The fragmentation protocol was modified to generate longer RNA fragments, with a median insert size of 200 bp. Briefly, after elution of the mRNA from binding beads (2 minutes at 80°C), sample processing continued immediately without further heat treatment. Indexed adapters were ligated onto cDNA fragments and the libraries were enriched with 13 cycles of PCR. The libraries were quantified, diluted and sequenced on a HiSeq2500 at the AGRF.

### DNA sequencing data, alignment and variant detection

DNA sequencing data from WEHI-AML-1 and WEHI-AML-2 (exome, genome and small amplicons) were aligned to the human genome (hg19) using bwa-mem v0.7.10-r789 with default settings^5^. Additional sequencing data were sourced from the NIH-NCI Genomic Data Commons (GDC) Data Portal [phs001027 and phs000178], through authorised access requests lodged through dbGaP. Aligned sequencing data (BAM files aligned to hg38) was downloaded from GDC, together with variant calls. Preliminary variant detection of WEHI-AML-1, WEHI-AML-2, EMC-AML-1, the EMC cohort [phs001027]^3^ and all TCGA-AML [phs000178]^6^ samples was performed with samtools version 0.1.19-44428cd, followed by Varscan v2.3.6^7^. These preliminary calls were refined with superFreq 0.9.15 with default settings, which also provided clonality estimates for copy number and somatic SNV calls^8^. Somatic mutation calls for SW1783 were available through the Catalogue of Somatic Mutations in Cancer (v81)^9^. The MutationalPatterns R Package was used to assess the trimer base context for somatic mutations^10^.

### Assessment of MBD4 status and proportion of CG>TG mutations in TCGA

To assess the frequency of CG>TG mutations in TCGA samples, we used somatic SNV calls available through GDC and applied a filter to restrict the analysis to variants detected with a variant allele frequency above 20%, with minimum 20 reads coverage and recognition by three out of the four callers: SomaticSniper, VarScan2, MuTect2, and MuSE. The consensus calling approach correlated well with results from our own analysis pipeline, both in terms of total mutational load and proportion of CG>TG mutations **(Extended Data Fig. 8)**.

Candidate germline loss-of-function variants impacting MBD4 were sourced from GDC and filtered to restrict the analysis to variants detected with a variant allele frequency above 10%, found with a population frequency below 1% in ExAC (non-TCGA cohort)^2^. The variant allele frequency and local copy number around *MBD4* were assessed in the matched cancer sample to designate cases as either monoallelic or biallelic inactivation.

### Assessment of the relative mutation rate and its association with genomic and epigenomic features

The incidence of CG>TG mutations was assessed in different genomic features, defined by the sequence context (NCG trimer, NCGN tetramer) or functional classification (CG island, promoter, promoter flanking, exon or intron). CG island annotation was obtained at the University of California, Santa Cruz (UCSC) genome browser^11^. Promoter and promoter flanking annotations were obtained from the Ensembl website through biomaRt^12^. Intronic and exonic regions were determined from the Ensembl v75 database^13^ (GRCh37).

As the mutations occurred almost exclusively in a CG context, the rate of CG>TG mutations per CG was calculated for each genomic feature. Each CG provides two mutational substrates and was therefore counted twice. Whole genome bisulfite (WGBS) data from CD34+ specimens from 5 healthy individuals was used to estimate methylation status for CG sites^14^. Due to the variable coverage in WGBS, the analysis was restricted to CG sites covered in at least 1 of the WGBS CD34+ samples, and a weighted average was used to estimate methylation status based on the coverage in each sample. The restriction to CG sites with coverage in WGBS ensured that a methylation call was available for all sites and omitted difficult to align regions.

A relative mutation rate (RMR) was calculated for each feature, either corrected for CG abundance (RMR–CG) or for 5mC abundance (RMR–5mC), using RoaMeR (an analysis script implemented in java)^15^. RMR is calculated in a way that is analogous to the fragments per kilobase of exon per million reads mapped statistic (FPKM). RMR-CG reflects the number of mutations expected in 1 Mb of CG sites in a particular genomic feature if 1000 somatic mutations were randomly selected from the total mutation load:

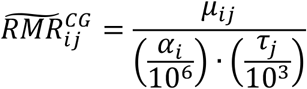

Where *μ*_*ij*_ represents the CG>TG mutation count for feature *i* within sample *j*, *α*_*i*_ represents the total number of CGs present in feature *i,* and *τ*_*j*_, is the total CG>TG mutation burden in sample *j*. The RMR-5mC replaces *α*_*i*_ with the weighted average methylation level for a feature:

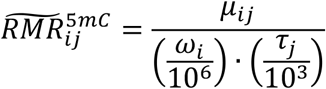

Where *ω*_*i*_ represents the sum of the weighted average methylation level for all CG sites in feature *i* across the CD34+ WGBS cases:

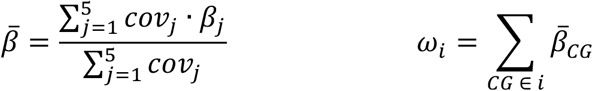

Where *coν*_*j*_ reflects the local coverage and *β*_*j*_ is the methylation level for this CG, both for sample _*j*_. For a fully methylated feature the CG-abundance (*α*_*i*_) is equal to the 5mCG-abundance (*ω*_*i*_), meaning the RMR-CG and RMR-5mC are equivalent.

RMRs were calculated for each genomic feature, then further stratified based on replication timing, transcriptional strand or expression level.

Replication timing: A conserved DNA replication timing profile was generated from 14 Repli-seq data sets from ENCODE^16^ downloaded through the UCSC genome browser^11^ (hg19). The data comprises genome-wide wavelet-smoothed values per 1kb bin for 14 cell lines (BG02ES, BJ, GM06990, GM12801, GM12812, GM12813, GM12878, Hela-S3, HepG2, HUVEC, IMR90, K562, MCF-7 and NHEK). The median value over all cell lines was calculated per 1-kb bin, then the median of these values was calculated for contiguous 10-kb blocks. Each 10-kb block was distributed among 4 equally-sized domains, designated the latest (< 29), late (≥ 29 & < 47), early (≥ 47 & < 63) and earliest (≥ 63) replication domains.

Transcriptional strand and expression level: Transcriptional strand bias analysis was performed by determining the template and non-template strands per gene as reported in Ensembl v75^13^. Overlapping gene bodies on the same strand were merged while overlapping regions defined by genes located on opposite strands were excluded. The number of CG>TG mutations on the template and non-template strands was counted genome-wide and corrected for the total CG-abundance within gene bodies. Transcriptional bias was assessed by binning genes into categories based on expression. The average FPKM value per gene was calculated from RNA-seq data available from WEHI-AML-1 and WEHI-AML-2. Genes with an average FPKM value ≤ 0.5 were considered to have very low expression (n=7464) and were allocated to the none-gene expression bin. The remaining genes were ordered by average FPKM value and distributed among 4 equally-sized expression bins (n=4509 per bin), designated lowest-, low-, high- and highest- gene expression bins. A high correlation between gene expression level and replication domain was noted.

RMR-5mC calculation for gnomAD: The Genome Aggregation Database^2^ (gnomAD) was provided by the Broad Institute and downloaded from Google Cloud Storage (release-170228). Rare CG>TG germline polymorphisms with a minor allele frequency between 0.0001 and 0.001 were considered for further analysis. SNPs were used in place of somatic mutations to calculate an RMR-5mC for all NCGN tetramers, either globally or for each replication domain.

### RNA sequencing alignment and splicing assessment

RNA-seq data from WEHI-AML-1 and WEHI-AML-2 was aligned with Tophat v2.0.12^17^. Aligned sequencing data for TCGA samples (BAM files aligned to hg38) was downloaded from GDC. Aligned RNA-seq data was counted over genes with featureCounts^18^. Reads spanning splice sites were used to calculate the proportion of reads involved in canonical and non-canonical splicing.

### Variant validation using a small amplicon panel

Primers flanking variants of interest were designed using Primer3 (http://bioinfo.ut.ee/primer3-0.4.0/). Whole genome amplification (WGA) was performed on DNA from WEHI-AML-1 and WEHI-AML-2 using the REPLI-g Mini/Midi Kit (Qiagen, Venlo, the Netherlands). Amplified DNA was purified using Agencourt AMPure XP (Beckman Coulter, Brea, CA, USA) before use in PCR, which was performed using Phusion Hot Start II High-Fidelity DNA Polymerase (Thermo Fisher Scientific, Waltham, MA, USA) (30 seconds at 98°C, 31 cycles of 10 seconds at 98°C, 20 seconds at 60°C and 15 seconds at 72°C, and a final 5 minutes at 72°C). PCR products were analysed on an agarose gel, pooled and purified using Agencourt AMPure XP. A second PCR was performed using indexed forward and reverse primers that introduce the P5 and P7 sequences (30 seconds at 98°C, 24 cycles of 10 seconds at 98°C, 20 seconds at 60°C and 15 seconds at 72°C, and a final 5 minutes at 72°C). PCR products were analysed on an Agilent 2200 Tapestation D1000 ScreenTape Assay (Agilent Technologies, Santa Clara, CA, USA), purified using Agencourt AMPure XP, quantified and diluted before sequencing on a MiSeq (Illumina, San Diego, CA, USA).

### Validation and phasing of WEHI-AML-1 *MBD4* variants

Primers flanking the *MBD4* variants in exon 3 and exon 7 were designed using Primer3. Non-amplified DNA extracted from remission material from WEHI-AML-1 was used as a template. PCR of exon 7 (MBD4_Ex7-F: 5’-CCTCTTGGTCTCTACGATCTTC-3’ and MBD4_Ex7-R: 5’-TCGGTAAGAGTCGTTGCCAT-3’) was performed using Platinum *Taq* DNA Polymerase (Thermo Fisher Scientific, Waltham, MA, USA) (5 minutes at 94°C, 35 cycles of 20 seconds at 94°C, 30 seconds at 60°C and 60 seconds at 72°C, and a final 5 minutes at 72°C). PCR of exon 3 (MBD4_Ex3-F: 5’-GCTGAAAGTGAACCTGTTGC-3’ and MBD4_Ex3-R: 5’-TGTGTTCTGAGTCTTTGGCTG-3’) was performed using Phusion Hot Start II High-Fidelity DNA Polymerase (30 seconds at 98°C, 32 cycles of 10 seconds at 98°C, 30 seconds at 60°C and 15 seconds at 72°C, and a final 10 minutes at 72°C). Phusion Hot Start II High-Fidelity DNA polymerase was used as its enhanced proof-reading ability was preferable for amplifying the poly-A stretch in exon 3. The PCR products were used directly for Sanger sequencing and analysed on Applied Biosystems 3730 or 3730*xl* capillary sequencers (Thermo Fisher Scientific, Waltham, MA, USA). Sanger sequencing results were analysed using SeqMan (DNASTAR, Madison, WI, USA) and by comparison to the human genome with BLAT (https://genome.ucsc.edu/cgi-bin/hgBlat). PCR products were also cloned using Zero Blunt TOPO PCR Cloning Kit and One Shot Top10 Electrocomp *E. coli* (Thermo Fisher Scientific, Waltham, MA, USA) prior to Sanger sequencing, to enable resolution of the mixed signal produced by frameshift variants. Plasmid DNA was obtained using PureLink HiPure Plasmid Miniprep Kit (Thermo Fisher Scientific, Waltham, MA, USA).

Total RNA from WEHI-AML-1 and WEHI-AML-2 at the time of first remission was used to make cDNA with SuperScript III Reverse Transcriptase (Thermo Fisher Scientific, Waltham, MA, USA). Primers flanking *MBD4* exons 3 to 8 were designed using Primer3 (MBD4_Phasing1-F 5’-GAGACCCTCAGTGTGACCAG-3’ with MBD4_Phasing1-R 5’-GCTGGAAAGGTGGTTGGTTG-3’, and MBD4_Phasing2-F 5’-GCTGAAAGTGAACCTGTTGC-3’ with MBD4_Phasing2-R 5’-GCTGGAAAGGTGGTTGGTTG-3’). PCR was performed using Phusion Hot Start II High-Fidelity DNA Polymerase (30 seconds at 98°C, 30 cycles of 10 seconds at 98°C, 30 seconds at 60°C and 2 minutes at 72°C, and a final 10 minutes at 72°C). The PCR products were cloned, plasmids were purified and Sanger sequenced, as above.

### Clonality analyses from bone marrow smears and single cells

EMC-AML-1 DNA was isolated from May-Grunwald Giemsa (MGG) stained bone marrow smears. Material was removed from glass slides and dissolved in RLT plus lysis buffer (Qiagen, Venlo, the Netherlands). Genomic DNA was isolated with the QIAsymphony. A Becton Dickson FACS Aria II (Franklin Lakes, NJ, USA) was used to sort intact and living cells into 96-well plates at one cell per well. WGA was performed on each cell using the REPLI-g Single Cell DNA Library Kit (Qiagen, Venlo, the Netherlands). The Illumina TruSight Myeloid Sequencing Panel (Illumina, San Diego, CA, USA) was applied to detect mutations in genes that are frequently altered in myeloid malignancies. Libraries were generated as per manufacturer’s instructions and the sequencing was performed on a MiSeq. Sanger sequencing was performed to assess mutations in *DNMT3A*, with the following PCR primers: R635-F: 5’-CAGGGTGTGTGGGTCTAGGA-3’ and R635-R: 5’-AAGCTTCCCCTTTGGGATAA-3’; R688-F: 5’-CAGGGAGATGGCTCCAAGTA-3’ and R688-R: 5’-TTTGCCCTTTACCCTCTCAA-3’; R882/A884-F: 5’-AGGAGTTGGTGGGTGTGAGT-3’ and R882/A884-R: 5’-TCTCCATCCTCATGTTCTTGG-3’; 3’UTR-F: 5’-TTCTAGAAGCCGCTGTTACCTC-3’ and 3’UTR-R: 5’-CCTCATCTAGCCCCCTTTTT-3’. PCR was performed using Taq DNA polymerase (Thermo Fisher Scientific, Waltham, MA, USA) (4 minutes at 94°C, 35 cycles of 1 minute at 94°C, 1 minute at 60°C and 1 minute at 72°C, and a final 7 minutes at 72°C). PCR products were purified using a Millipore Microscreen™ PCR cleanup plate (Merck, Kenilworth, NJ, USA) and sequenced on an Applied Biosystems 3130xl Genetic Analyzer (Thermo Fisher Scientific, Waltham, MA, USA).

### Site-directed mutagenesis and cloning

Full length *DNMT3A* and *MBD4* cDNA were PCR amplified from a pReceiver-B13 vector (GeneCopoeia, Rockville MD, USA) and cloned into pET28-MHL expression vector (GenBank accession EF456735, Addgene, Cambridge, MA, USA) by applying BD-BioScience In-Fusion (Franklin Lakes, NJ, USA) enzyme-mediated directional recombination between complementary 15 nucleotide DNA sequences at the end of the PCR products, using the following primers for DNMT3A: sense 5’-TTGTATTTCCAGGGCCCCGCCATGCCCTCCAGC-3’ and anti-sense 5’-CAAGCTTCGTCATCACACACACGCAAAATACTCCTTCAGC-3’, MBD4: sense 5’-TTGTATTTCCAGGGCGGCACGACTGGGCTGGAGAGTCT and anti-sense 5’-TTGTATTTCCAGGGCGGCACGACTGGGCTGGAGAGTCT-3’. The truncated form of MBD4, residues 430-580, in the pET28 backbone was sourced from Addgene.

QuikChange II XL Site-Directed Mutagenesis Kit (Agilent Technologies, Santa Clara, CA, USA) was used to generate the DNMT3A and MBD4 mutants. The primer sequences were as follows, with the site of deletion or mutation underlined: MBD4 H567 deletion: sense 5’-CCCTGAAGACCACAAATTAAATAAATAT___GACTGGCTTTGGGAA-3’ and anti-sense 5’-TTCCCAAAGCCAGTC___ATATTTATTTAATTTGTGGTCTTCAGGG-3’; MBD4 D560A: sense 5’-GAAGCAGGTGCACCCTGAAGCCCACAAATTAAATAAATATCA-3’ and anti-sense 5’-TGATATTTATTTAATTTGTGGGCTTCAGGGTGCACCTGCTTC-3’; DNMT3A R635W: sense 5’-AGAGGAAGCCCATCTGGGTGCTGTCTCTC-3’ and anti-sense 5’-GAGAGACAGCACCCAGATGGGCTTCCTCT-3’; DNMT3A R688C: sense 5’-GTCGGGGACGTCTGCAGCGTCACAC-3’ and anti-sense 5’-GTGTGACGCTGCAGACGTCCCCGAC-5’; DNMT3A R882C: sense 5’-TCTCCAACATGAGCTGCTTGGCGAGGCAG-3’ and anti-sense 5’-CTGCCTCGCCAAGCAGCTCATGTTGGAGA-3’; and DNMT3A A884V: sense 5’-CATGAGCCGCTTGGTGAGGCAGAGACTGC-3’ and anti-sense 5’-GCAGTCTCTGCCTCACCAAGCGGCTCATG-3’. Successful mutagenesis was confirmed by enzymatic digestion and DNA sequencing.

### Expression and purification of recombinant DNMT3A, MBD4 and TDG proteins

The pET28-MHL containing hexahistidine-tagged DNMT3A wild-type and mutants, MBD4 full length and mutants, MBD4 residues 430-580 and mutants, and TDG residues 111-348 (kindly provided by Hideharu Hashimoto and Xiaodong Cheng) were expressed in *Escherichia coli* BL21(DE3) Gold and BL21(DE3) pLysS cells (Thermo Fisher Scientific, Waltham, MA, USA). A single BL21(DE3) colony containing the pET28 expression vector with the desired insert was cultured for 16 hours at 37°C in Luria Broth (LB) medium supplemented with 50 µg/ml kanamycin. The culture was diluted 25 times in LB medium without selection marker and grown at 37°C until it reached an OD600 of 0.5 to 0.8. Cells were induced by adding 0.5 mM isopropyl b-D-1-thiogalactopyranoside and incubated for 4 hours at 37°C. Cell cultures were centrifuged at 38,000 × g for 30 minutes at 4°C and the cell pellets resuspended in lysis buffer containing 20 mM sodium phosphate, 500 mM NaCl, 10 mM imidazole (pH 7.4), 1 mg/ml lysozyme, 200 µg/ml DNAse, and 1x SIGMA*FAST* Protease Inhibitor, EDTA-Free (Sigma-Aldrich, St. Louis, MO, USA), before incubation on ice for 30 minutes. The cells were sonicated on ice for 6 minutes at amplitude 60%, using 10 second bursts followed by a 20 second reprieve with a Branson Digital Sonifier (Branson, Danbury, CT, USA). The lysate was clarified by centrifugation at 38,000 × g for 30 minutes at 4°C. The hexahistidine-tagged proteins were isolated from the crude lysate using Ni-NTA Superflow (Qiagen, Venlo, the Netherlands) and a 2.5 × 10cm Econo-chromatography column (Bio-Rad, Hercules, CA, USA). The Ni-NTA resin was washed twice with cold washing buffer containing 20 mM sodium phosphate, 500 mM NaCl and 10 mM imidazole (pH 7.4) to remove non-specific interacting proteins. Hexahistidine-tagged proteins were eluted from the Ni-NTA group on the matrix with cold elution buffer, containing 20 mM sodium phosphate, 500 mM NaCl and 500 mM imidazole (pH 7.4). Directly after elution into buffer containing 500 mM imidazole, the protein suspensions were dialyzed in a Slide-A-Lyzer G2 Dialysis Cassette (gamma irradiated, 10K MWCO) (Thermo Fisher Scientific, Waltham, MA, USA), for 2 hours at 4°C against 300x volume dialysis buffer containing 50 mM Tris-HCl and 150 mM NaCl (pH 7.6). The dialysis buffer was refreshed twice, with further incubation at 4°C for 2 hours and 16 hours. Proteins were quantified using Qubit protein assay kit and Qubit 3.0 fluorometer (Thermo Fisher Scientific, Waltham, MA, USA). Proteins were verified by SDS-PAGE using a NuPage Novex 4-12% Bis-Tris Protein Gel run in a Bis-Tris XCell SureLock™ Mini-Cell system (Thermo Fisher Scientific, Waltham, MA, USA) with 1x MOPS at 200V for 90 minutes. Blots were incubated in blocking buffer containing 5% BSA, 0.1% Tween and 1xPBS with the appropriate antibodies: α-His H-15 (sc-803, Santa Cruz Biotechnology, Dallas, TX, USA), α-DNMT3A (ab2850, Abcam, Cambridge, UK), α-TDG (ab106301, Abcam), and α-MBD4 (ab12187, Abcam). Proteins were visualized using a Li-Cor Odyssey 3.0 (Li-Cor, Lincoln, NE, USA).

### MBD4 and TDG glycosylase activity assays

MBD4 and TDG glycosylase activity assays were performed as described (Hashimoto et al., NAR, 2012). Briefly, MBD4 and TDG glycosylase activity was measured with 32 bp oligonucleotides labelled with 6-carboxy-fluorescein (FAM) and G-T mismatch excision was monitored by denaturing gel electrophoresis following NaOH hydrolysis. The 32 bp oligonucleotides used were (FAM)-5′-TCGGATGTTGTGGGTCAGXGCATGATAGTGTA-3′ (where X = C or T) and 5′-TACACTATCATGCGCTGACCCACAACATCCGA-3′ (Integrated DNA Technologies, Coralville, IA, USA). Double-stranded FAM labelled match and mismatch substrates were prepared by hybridisation. 100 µM of oligodinucleotides were mixed in 50 µl annealing buffer containing 10 mM Tris HCl, 1 mM EDTA and 50 mM NaCl (pH 8.0), then incubated at 95° C for 2 minutes, followed by a steady temperature reduction over 45 minutes to 25° C. The double-stranded duplexes were cooled and stored at 4°C. MBD4 (500 nM), TDG (40 nM) and various concentrations of DNMT3A protein were mixed with 100 nM double-stranded FAM labelled 32 bp substrates in 20 µl nicking buffer (10 mM Tris–HCl, pH 8, 1mM EDTA and 0.1% BSA) and incubated at 37°C for 60 minutes. Reactions were terminated by adding 2 µl 1 M NaOH, before boiling for 10 minutes, prior to addition of 20 µl of loading buffer (98% formamide, 1 mM EDTA and 1 mg/ml of Bromophenol Blue and Xylene Cyanole). The samples were boiled for 10 minutes, immediately cooled on ice and then loaded onto a 8.3 cm × 7.3 cm, 1 mm thick denaturing gel containing 7 M urea, 24% formamide, 15% acrylamide and 1x Novex TBE Running Buffer (Thermo Fisher Scientific, Waltham, MA, USA). The gels were run on a Mini-PROTEAN Tetra Cell (Bio-Rad, California, USA) in 1× TBE buffer at 100 V for 90 minutes. FAM-labeled single-stranded DNA was visualised and images acquired using the 473nm laser (Blue LD Laser) and 530DF20 emission filter on a Typhoon FLA9500 (GE Healthcare, Illinois, USA).

### Data availability

Sequencing data from WEHI-AML-1 and WEHI-AML-2 have been deposited at the European Genome Phenome Archive (EGA) [EGAS00001002581]. The data are available for ethically approved research into haematological malignancy upon completion of a data transfer agreement. Sequencing data from EMC-AML-1 were sourced from the dbGaP under accession phs001027. TCGA data were downloaded from the GDC Data Commons. Code to reproduce the figures and data are made available through GitHub (https://github.com/MathijsSanders/AML-RoaMeR).

